# From group-level neurophysiological biomarkers to individual-level phenotyping to identify mechanisms of neurodegenerative disease

**DOI:** 10.64898/2026.10.26.761866

**Authors:** Amirhossein Jafarian, Laura E Hughes, Michelle Naessens, Matthew A. Rouse, Thomas E Cope, Negin Holland, Alexander Murley, Sean Tan Yong Wei, Timothy Rittman, David J. Whiteside, James B Rowe

## Abstract

Precision medicine approaches require physiological effects to be informative when translated to the individual. Reliable group differences may have limited individual value when distributions overlap, mechanisms are heterogeneous, or patients differ in phenotype. A feature can differ between group means and still overlap heavily across individuals Moreover, combinations of physiological abnormalities may reveal subject-level structure that is not captured by any single marker at the group level. We evaluate the translation between group- and individual-level features derived from resting-state magnetoencephalography (MEG) using behavioural-variant frontotemporal dementia (bvFTD) and progressive supranuclear palsy (PSP) as demonstrator conditions. These illnesses have distinct molecular pathology but share the pressing need for therapeutic development and have partially overlapping clinical and physiological features. We combine clinical phenotype, power spectral density (PSD), and latent network mechanisms inferred by dynamic causal modelling (DCM) to compare groups and to access their individuals mapping translations. We used classification as an exploratory probe of how group-defined physiological features from 74 healthy controls, 45 participants with bvFTD and 55 participants with PSP support individual assignment. Clinical scores showed strong individual-level translation, consistent with clinical diagnostic definitions (e.g, controls-vs-PSP accuracy 0.94; bvFTD-vs-PSP accuracy 0.83). MEG spectra feature classified individuals weakly (accuracy ∼0.6), despite group level re-organization (e.g., reduced beta-band and increased fronto-insular alpha-phase/beta-amplitude coupling in bvFTD). Inferred biophysical parameters gave intermediate performance (e.g. PSP-vs-control accuracy 0.70; bvFTD-vs-PSP accuracy 0.74) while retaining mechanistic interpretability. Using this approach, experimental medicine studies of neurological disease can formally trade between mechanistic insight and classification performance, in using biomarkers that can account for individual heterogeneity within and between diagnostic groups.

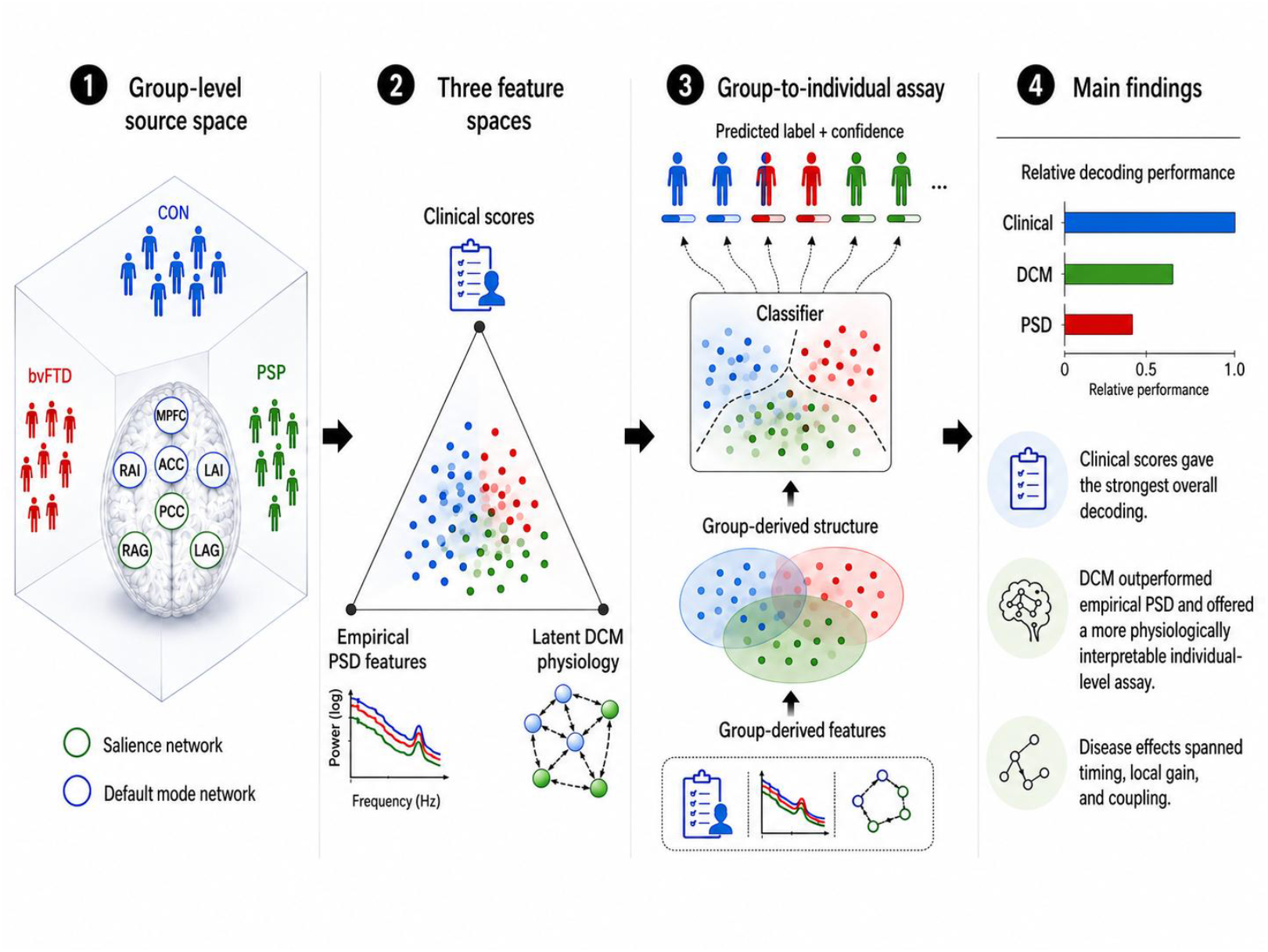

## Introduction

There is a pressing need for quantitative biological tools for *in vivo* human use that improve mechanistic understanding of neurodegenerative disease and remain interpretable at individual level. Clinical assessments measure the cognitive, behavioural, and motor consequences of pathology and remain the foundation of diagnosis, but they do not explain the processes generating these deficits. In contrast, neuroimaging and other biomarkers can reveal molecular, structural, or network-level changes, or latent causes of neurophysiological changes. Biomarker-based tests of mechanistic hypotheses are particularly important when similar clinical presentations arise from distinct neural dynamics, However, biomarkers may compromise clinical utility for mechanistic interpretability; or be reliable only at the group level rather than for individuals.

We examined the tradeoff between mechanistic insights and classification performance (cf. diagnosis), in the context of behavioural-variant frontotemporal dementia (bvFTD) and progressive supranuclear palsy (PSP). These conditions overlap clinically, with shared features including apathy, disinhibition, impaired social cognition, executive dysfunction, and parkinsonism, despite distinct molecular pathologies (Shi et al. 2021, Murley et al., 2020). Indeed, PSP pathology can underlie a bvFTD presentation or PSP–Richardson’s syndrome (PSP-RS) as the disease progresses. bvFTD is associated with degeneration of fronto-insular, anterior cingulate, and orbitofrontal cortex, with underlying FTLD-tau or FTLD-TDP pathology (Rascovsky et al., 2011; Murley et al., 2020; Seeley et al., 2009). PSP is a 4R-tauopathy with distributed subcortical and cortical pathology (Höglinger et al., 2017; Kovacs et al., 2020). At the group level, bvFTD and PSP both disrupt the macroscopic salience-network, with reciprocal changes in the default-mode network (Seeley et al., 2007; Seeley et al., 2009; Zhou et al., 2010; Chiong et al., 2013; Whitwell et al., 2011; Rittman et al., 2016; Cope et al., 2018; Bharti et al., 2017; Brown et al., 2017). The anterior insula and anterior cingulate that form the salience-network, are involved in detecting behaviourally relevant events and regulating transitions between internally and externally oriented control systems (Sridharan et al., 2008; Menon and Uddin, 2010; Goulden et al., 2014). However, group level physiological differences are not necessarily sufficient for individualised phenotyping. A feature may differ significantly between groups yet have limited value for one participant when data distributions overlap, broad, or covary. Conversely, several weak physiological effects may become individually informative when combined. This distinction matters in heterogeneous neurodegenerative disorders, where distributed reorganization and divergent physiological perturbations may induce similar symptoms.

We sought to distinguish three issues that are often conflated in biomarker studies: (i) whether a feature differs between groups; (ii) whether that feature, or a combination of features, remains meaningful at the level of individual participants; and (iii) what the pattern of individual-level differences reveals about disease mechanisms.

First, we characterised clinical, spectral, and latent pathophysiological features using strength of evidence, effect magnitude, combined evidence-effect scores, and recurrence across contrasts. Second, classification used exploratory group-to-individual assays conditional on full-sample feature selection, rather than diagnostic validation. Participants were excluded from classifier fitting, not feature selection. This follows the category theory approach, with potential mappings between inferential domains. Group-level feature representations are transformed through statistical comparison, ranking and panel selection. The resulting representation and decision rules are then applied to an individual. This translation preserves the order and composition of the analytical mappings. Classification accuracy is descriptive, not independent evidence of individual-level translation. This approach is relevant to precision medicine because individualized treatment, stratification, and monitoring require more than ‘significant’ group differences. They require features that are interpretable, physiologically meaningful, and coherent at the subject-level. However, there may be a trade-off between classification performance and mechanistic insights.

In this study, we therefore compare (i) clinical scores, that explicitly refer to syndromic expression; (ii) empirical spectral features from magnetoencephalography, that quantify neuronal dynamics; and (iii) physiological parameters inferred using DCM, to quantify the effects of disease on meso- and macro-scale networks that generate the magnetoencephalography signals. In doing so, we bridge between conventional case-control neuroimaging methods and individualised physiological phenotyping.

## Results

A glossary of abbreviation is provided in the Appendix A. A summary of clinical and demographic information on participants is given in Table 1. Further clinical information, network information, signal features, and classifier terms are presented in supplementary information sections A, B, C and D. The summary of frequency analysis is illustrated in Figure 1. To align the results with the clinical and physiological questions, findings are reported contrast-by-contrast and a summary presented in Figure 2.

**Table 1.** Demographic and clinical characteristics, and their comparison between groups by ANOVA or chi-squared tests as appropriate. ACE-R, revised Addenbrookes Cognitive Examination, MMSE mini-mental state examination, CBI-R revised Cambridge Behavioural Inventory, PSPRS Progressive supranuclear palsy rating scale.

|  | **CONTROL** | **bvFTD** | **PSP** | **p-value** |
| --- | --- | --- | --- | --- |
| N | 74 | 45 | 55 | — |
| Age, mean ± SD | 65.5 ± 7.5 | 63.3 ± 6.9 | 67.7 ± 7.5 | 0.011 |
| Age range, years | 45–85 | 49–80 | 49–83 | — |
| Sex, female/male | 31/43 | 13/32 | 21/34 | 0.360 |
| ACE-R (0-100) | 96.00 | 63.69 | 79.75 | <0.001 |
| MMSE (0-30) | 29.22 | 21.90 | 26.84 | <0.001 |
| CBI-R total (0-180) | — | 87.71 | 55.49 | <0.001 |
| PSPRS (0-100) | — | 13.80 | 36.69 | <0.001 |

**Figure 1.**
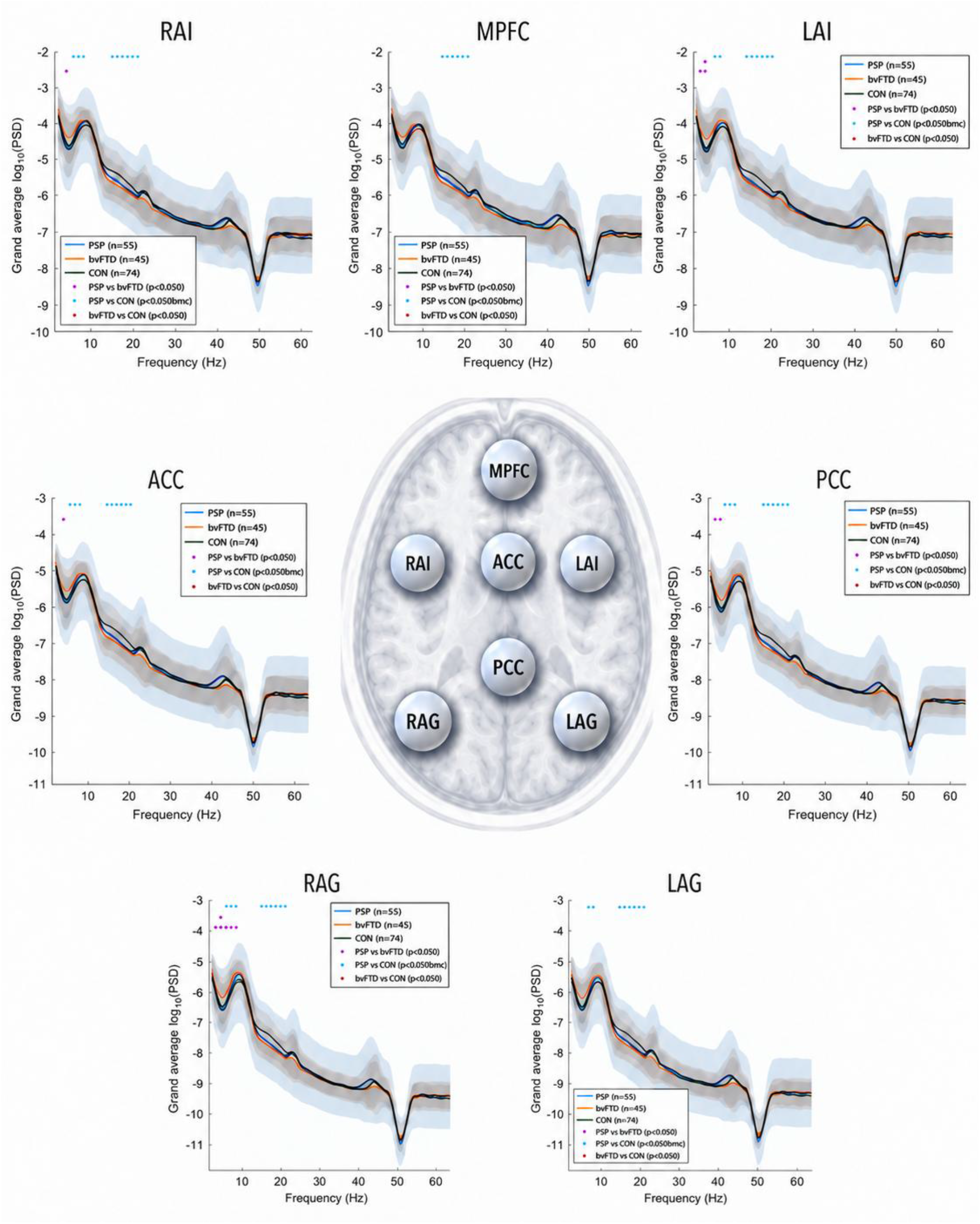
Source-space resting-state spectral responses across the salience/default-mode network in PSP, bvFTD, and controls. Group-averaged source-space log-transformed power spectral density (log10 PSD) is shown for the seven MEG source regions: left anterior insula (LAI) and right anterior insula (RAI), medial prefrontal cortex (MPFC), anterior cingulate cortex (ACC), posterior cingulate cortex (PCC), right angular gyrus (RAG), and left angular gyrus (LAG). Source time courses were extracted after participant-specific MRI-informed source inversion, and PSD was estimated from resting-state MEG using Welch’s method and visualised as log10 PSD across the frequency range. Blue, orange, and green traces show PSP, bvFTD, and control groups, respectively, with shaded regions indicating group variability. Coloured dots mark frequency bins showing nominal pairwise group differences: purple for PSP-bvFTD, magenta for PSP-vs-controls, and cyan for bvFTD-vs-controls. The central panel provides a schematic anatomical spatial reference for the source locations.

**Figure 2.**
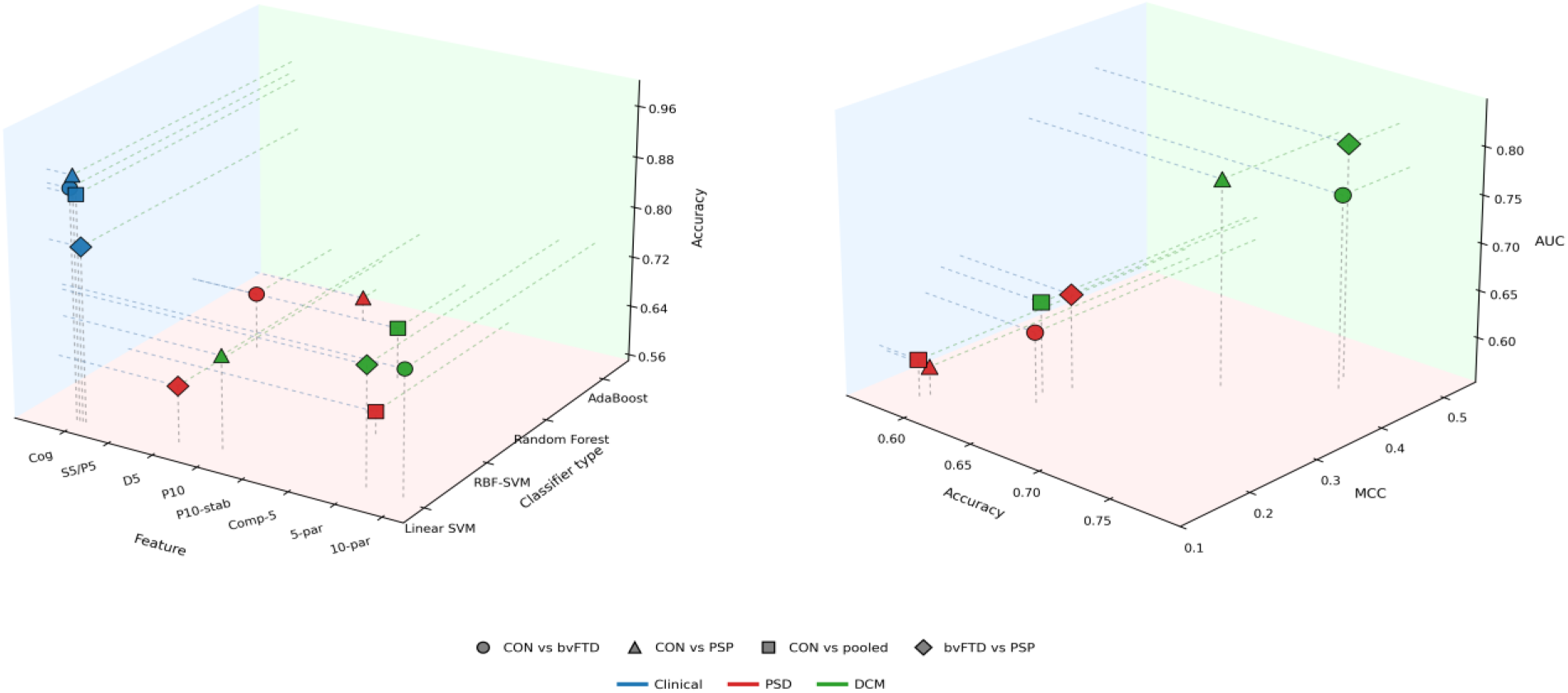
Group-to-individual performance using clinical, PSD, and DCM features. The displayed results are exploratory probe using classification and conditional on the full-sample feature selection. The left panel shows best accuracy as a function of feature type and classifier family. Clinical/cognitive-scores provide the strongest individual-level coherence (as expected from clinical diagnostic criteria). In contrast, PSD-based models are less accurate, indicating that empirical spectral abnormalities, although physiologically informative, translate only modestly to subject-level assignment. DCM provide intermediate accuracy, suggesting that latent parameters provide a more coherent subject-level representation than empirical spectral features alone. The right panel shows PSD and DCM according to Matthew’s correlation coefficient (MCC) and Accuracy (AUC), where accuracy alone is not sufficient to indicate the physiological usefulness of a model. The separation between PSD and DCM is consistent with a more coherent latent physiological representation in this selected sample but does not establish an independently validated advantage in generalisation.

### Controls versus bvFTD

The clinical data showed the clearest overall patient-control separation. Clinically, ACE-R, MMSE, and FAB showed strong deficits in bvFTD. In the group-to-individual assay, ACE-R alone achieved an accuracy of 0.913 and FAB achieved 0.869, while the multivariate clinical model reached 0.915. This indicates that the behavioural-cognitive burden measured by these scales is not only a group-level difference but a highly coherent individual-level expression of bvFTD.

Frequency-resolved statistics showed spatially extensive differences across both anterior salience-related and posterior/default-mode sources, with significant bins concentrated around approximately 5-7 Hz and 14-20 Hz. bvFTD spectra showed relatively greater low-frequency power and relatively reduced upper-alpha/low-beta power. Physiologically, this pattern suggests organised fast rhythms are weaker, and slower components occupy a larger proportion of the spectrum. The feature analysis converged on this interpretation: 20 PSD features survived Benjamini-Hochberg correction, dominated by increased within-source alpha-phase/beta-amplitude coupling (PAC) and reduced beta residual maximum across all seven channels, with additional reductions in relative beta power and spectral entropy in a subset of regions.

In PSD-feature prediction, the highest AUC was achieved by a random forest using the P5 panel, comprising PAC in RAI and LAI, beta residual maximum in RAI and LAI, and relative beta power in RAI (AUC 0.627; accuracy 0.597). The best overall operating point was obtained by random forest, using the S5 panel of PAC and beta-residual features in RAI, LAI, and ACC (accuracy 0.639; F1-score 0.494; MCC 0.215; AUC 0.617). Note how the clear group-level bvFTD spectral phenotype translates only modestly to individual assignment.

DCM-based prediction was stronger than PSD prediction. A linear SVM using a ten-parameter panel 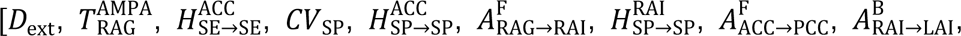, 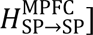 achieved accuracy 0.748, specificity 0.878, MCC 0.446, and AUC 0.756. The selected parameters were diverse, combining extrinsic transmission delay [*D*_ext_], right angular-gyrus AMPA-mediated excitatory integration 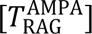, superficial-pyramidal membrane capacitance [*CV*_SP_], spiny-excitatory self-coupling in ACC 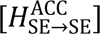, superficial-pyramidal self-coupling in ACC, RAI, and MPFC 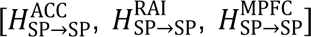, forward AMPA coupling from RAG to RAI and from ACC to PCC 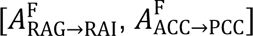, and backward AMPA coupling from RAI to LAI 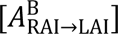. The implication is that bvFTD-control separation may be better expressed as a distributed alteration in cortical timing, local excitatory/superficial-pyramidal gain, and AMPA-mediated SN-DMN communication than as a single local abnormality. Thus, bvFTD produced group-level spectral and physiological effects that retained moderate subject-level coherence.

### Controls versus PSP

PSP was clearly separated from controls clinically in the clinical analysis. Compared with controls, PSP participants had lower ACE-R scores (79.75 versus 96.00; Cohen’s d=-2.305), lower MMSE scores (26.84 versus 29.22; d=-1.232), and lower FAB scores (11.82 versus 17.25; d=-2.084), all surviving Benjamini-Hochberg FDR correction. In the group-to-individual assay, the multivariate clinical model achieved accuracy 0.936, while ACE-R alone achieved accuracy 0.904, FAB 0.863, and MMSE 0.775. These confirm that PSP-control separation is not only clinically evident at the group level but is also strongly expressed at the individual level across global cognitive and frontal-executive measures.

Frequency-resolved statistics showed focal and low-frequency effects of PSP rather than spatially extensive changes: LAI showed significant bins at approximately 2.0 and 3.0 Hz, RAG showed a significant bin at approximately 3.0 Hz, and ACC, RAI, MPFC, PCC, and LAG showed no significant bins in this comparison. No PSD feature survived Benjamini-Hochberg FDR correction. The leading uncorrected trends involved flatter or less negative 1/f slopes in PSP across sources, higher beta peak frequency in PSP, and smaller focal contributions from relative gamma in LAG, alpha residual/relative alpha in PCC, and relative delta in LAI. This suggests that PSP does not produce the same broad spectral slowing and beta disorganisation as bvFTD, despite marked clinical abnormalities. The highest PSD classification result was obtained with AdaBoost using the D5 panel, comprising 1/f slope in ACC, MPFC, LAG, and RAG plus relative gamma power in LAG. This model achieved accuracy 0.589, sensitivity 0.509, specificity 0.649, precision 0.519, F1-score 0.514, MCC 0.158, and AUC 0.582 (TP 28, TN 48, FP 26, FN 27). The CON-PSP PSD effects are weak, focal, and poorly coherent at the individual level.

DCM prediction yielded moderate accuracy where the best result was obtained by a linear SVM using the P10 panel 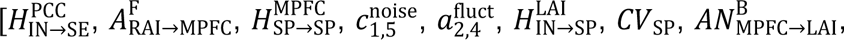 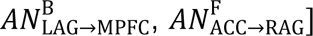, with accuracy 0.698, sensitivity 0.491, specificity 0.851, precision 0.711, F1-score 0.581, MCC 0.371, and AUC 0.770 (TP 27, TN 63, FP 11, FN 28; N=129). In physiological terms, the selected panel combined local microcircuit parameters, capacitance, receptor-specific coupling, and modelled variability: inhibitory-to-spiny-excitatory coupling in PCC 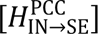, forward AMPA coupling from RAI to MPFC 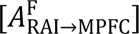, superficial-pyramidal self-coupling in MPFC 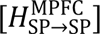, inhibitory-to-superficial-pyramidal coupling in LAI 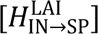, superficial-pyramidal capacitance [*CV*_SP_], backward NMDA coupling from MPFC to LAI 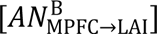, backward NMDA coupling from LAG to MPFC 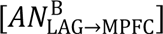, forward NMDA coupling from ACC to RAG 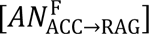, and two modelled variability/noise terms [specific sensor noise 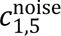 and endogenous fluctuation 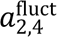]. Thus, PSP-vs-control separation was clinically expected, spectrally focal, and moderate in DCM space.

### Controls versus the combined patient group

Pooling bvFTD and PSP against controls preserved a clear disease-control difference clinically but did not strengthen individual-level physiological prediction. Single-feature prediction remained high for ACE-R (0.907) and FAB (0.855), whereas the multivariate clinical model reached 0.835. This indicates that the pooled patient-control contrast is dominated by a shared axis of global cognitive and frontal dysfunction rather than by complementary information unique to each diagnosis.

No PSD derived feature survived Benjamini-Hochberg FDR correction in the CON-pooled-patient contrast (105 features; 5 uncorrected p<0.05; no p<0.01 effects; minimum p=0.029; minimum q=0.446). The leading uncorrected features were beta residual maximum in RAI, RAG, MPFC, ACC, LAG, PCC, and LAI, relative theta power in RAI, and alpha residual maximum in LAG and PCC. Controls showed higher beta residual and alpha residual structure, whereas pooled patients showed higher RAI theta. Physiologically, this suggests a shallow common disease axis involving weaker organised beta/alpha rhythmic structure and a modest slowing component, but not a corrected pooled-patient PSD signature. In the group-to-individual assay, control-pooled-patient PSD features-based classification is weak. The strongest threshold-independent PSD result was obtained using the compact p-value/effect-size-ranked five-feature panel with an RBF-SVM, comprising beta residual maximum in RAI, RAG, MPFC, and ACC plus relative theta power in RAI. This model achieved accuracy 0.586, sensitivity 0.660, specificity 0.486, precision 0.635, F1-score 0.647, MCC 0.148, and AUC 0.591 (TP 66, TN 36, FP 38, FN 34; N=174). A sensitivity-biased AdaBoost solution with the same five-feature panel reached higher apparent accuracy (0.621) and sensitivity (0.880), but with low specificity (0.270) and poor AUC (0.476), thereby it was not treated as stronger physiological evidence. In summary, pooling syndromes preserve weak shared spectral structure but reduces the individual-level coherence of syndrome-specific PSD features.

In the DCM statistical analysis, no single parameter survived FDR correction (minimum p=0.00282; minimum q=0.410). The strongest classification operating point was obtained with a Random Forest using the P10/stability ten-parameter panel, with accuracy 0.632, sensitivity 0.720, specificity 0.514, precision 0.667, F1-score 0.692, MCC 0.238, and AUC 0.639 (TP 72, TN 38, FP 36, FN 28; N=174). The highest threshold-independent AUC was obtained by a linear SVM using the compact five-parameter panel 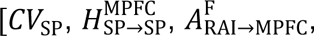 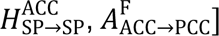 (AUC 0.648; accuracy 0.598; MCC 0.174). The ten-parameter DCM panels were dominated by superficial-pyramidal capacitance and self-coupling 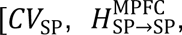 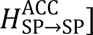 directed AMPA communication linking RAI, MPFC, ACC, PCC, and RAG 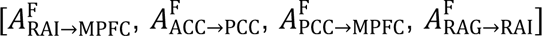, ACC spiny-excitatory self-coupling 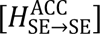, LAI inhibitory timing 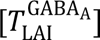, and a specific sensor-noise term 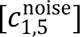. This is consistent with a shallow common disease-control axis involving local excitability/gain and effective communication across the SN-DMN network, but the weak-to-modest AUC and MCC indicate that combining bvFTD and PSP dilutes syndrome-specific physiological structure.

### bvFTD versus PSP

Direct comparison of the two syndromes confirmed a clinical dissociation between behavioural-cognitive and motor phenotypes. PSP showed higher PSPRS burden than bvFTD (36.69 versus 13.80; Cohen’s d=-1.880 for bvFTD minus PSP; FDR-corrected p=3.81e-11), and PSPRS alone achieved classification accuracy 0.843. CBI-R showed higher values in bvFTD than PSP (87.71 versus 55.49; d=1.016; FDR-corrected p=1.19e-4) and single-score accuracy 0.702. ACE-R and MMSE were lower in bvFTD than PSP (ACE-R: 63.69 versus 79.75, d=-0.908; MMSE: 21.90 versus 26.84, d=-0.893), both surviving FDR correction and achieving single-score accuracies of 0.688 and 0.691, respectively. The multivariate clinical model achieved accuracy 0.828 confirming syndrome separation driven primarily by PSP motor-functional burden, with additional behavioural and cognitive contributions.

In the frequency domain, differences were focal and concentrated at low frequencies. LAI showed differences 2.0 and 3.0 Hz, ACC at approximately 3.0 Hz, PCC at approximately 2.0 and 3.0 Hz, and RAG at approximately 2.0, 3.0, 5.0, 6.0, and 7.0 Hz; RAI, MPFC, and LAG showed no significant effects. 1/f slope survived FDR across all seven SN-DMN sources, and relative beta survived FDR in LAI and RAI, with trend-level beta effects in other regions. Thus, the frequency-bin analysis supports a focal low-frequency syndrome difference, whereas the derived-feature analysis supports a broader aperiodic/beta organisation difference. The derived-feature analysis indicated more negative 1/f slopes in bvFTD than PSP participants across sources, together with lower relative beta in bvFTD in key salience-related regions. A more negative 1/f slope across sources indicates broader aperiodic redistribution, which may reflect altered asynchronous population activity, changes in synaptic balance, or altered excitatory-inhibitory organisation at the field-potential level. In contrast, the frequency-bin comparison shows that narrowband differences between PSP and bvFTD in the raw logPSD spectra are focal and low-frequency, especially in LAI, PCC, and RAG. This distinction is important because it avoids treating the derived 1/f/beta feature pattern as if it were identical to the frequency-resolved result. In the PSD classification, the highest AUC was obtained by a linear SVM using ten-feature panels dominated by 1/f slope across all seven sources and relative beta in RAI, LAI, and RAG (AUC 0.652; accuracy 0.610; sensitivity 0.533; specificity 0.673; F1-score 0.552; MCC 0.208). The strongest accuracy and MCC were obtained by AdaBoost using the D5 panel (accuracy 0.640; sensitivity 0.533; specificity 0.727; precision 0.615; F1-score 0.571; MCC 0.266; AUC 0.621).

The DCM classification analysis supported stronger subject-level separation. The best DCM operating point was a linear SVM using a compact five-parameter panel 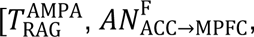 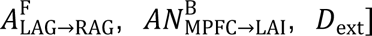, with accuracy 0.740, sensitivity 0.800, specificity 0.667, precision 0.746, F1-score 0.772, MCC 0.472, and AUC 0.798 (TP 44, TN 30, FP 15, FN 11). The five selected DCM parameters point to the RAG AMPA-mediated excitatory time constant 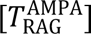, forward NMDA coupling from ACC to MPFC [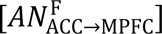, forward AMPA coupling from LAG to RAG 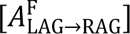, backward NMDA coupling from MPFC to LAI 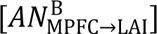, and extrinsic conduction/transmission delay [*D*_ext_].

## Discussion

This study sought to determine whether clinical, spectral, or latent-neurophysiological features of brain function differ not only at the group level but are sufficiently consistent to translate into individual-level classification. We confirm that magnetoencephalography can reveal cortical mechanisms of disease, but at a cost in terms of heuristic performance for participant classification. The biophysical modelling of pathophysiology, to the cellular and synaptic level, can provide a bridge to animal models and pharmacological interventions (Moran et al., 2011). The latent physiological parameters inferred from biophysical models capture non-linear, non-trivial features that provide more accurate classification than power spectral features extracted from the same non-invasive imaging data. However, this advantage for understanding disease mechanisms comes with a reduction in classification performance (c.f. clinical diagnostics), compared to clinical data.

Neurological disease may be expressed through a few dominant features or through combinations of individually modest changes. Alongside approaches centred on individual biomarkers, such as neurofilament light or pTau217, our framework examines how group-defined feature combinations support individual-level characterisation. Clinical scores showed the strongest group-to-individual correspondence in this cohort, consistent with the role of symptoms and signs in clinical diagnosis. High classification accuracy has also been reported using caregiver observations of behavioural changes (Murley et al., 2025). These measures characterise the clinical expression of disease but do not directly quantify the synaptic and network processes underlying disease expression.

Empirical spectral feature panels supported weaker individual assignment, including in contrast with significant group-level effects. DCM-derived parameter panels, by comparison, supported moderate individual assignment while providing model-based interpretations in terms of synaptic timing, local gain and network coupling. Their contribution therefore extends beyond classification accuracy; they connect the observed organisation of neuronal activity to candidate physiological explanations. This contrast illustrates why group-level statistical prominence, individual-level discriminability and mechanistic interpretability should be examined together, rather than treating anyone as a sufficient indicator of a feature’s scientific value.

By applying complementary classifiers as probes of panels, we characterise how group-prioritised features support individual assignment within the study cohort. The methodological contribution is a structured examination of how feature representation, panel composition and decision mapping shape this correspondence. This gives multivariate MEG analysis a complementary role alongside clinical assessment in which characterising physiological patterns and generating mechanistic hypotheses relevant to individual phenotyping. More broadly, the framework provides a way to interrogate the route from group findings to individuals across biomarker studies, while identifying candidate representations for independent and longitudinal investigation in experimental medicine.

Time-frequency analyses of bvFTD and PSP have previously identified reduced beta power and altered fronto-insular cross-frequency coordination (Hughes et al., 2018; Sami et al., 2018; Zhang et al., 2016). Resting-state activity is shifted toward slower cortical dynamics, with reduced network efficiency and impaired maintenance of organised fast states (Azami et al., 2023; Baik et al., 2022; Bruna et al., 2023). Reduced beta residual power, after accounting for the aperiodic background, indicates a reduction in organised beta-band rhythmic structure over and above a broad spectral tilt. Such changes in beta-band dynamics and cross-frequency coupling reflect behavioural and functional decline (Hughes et al., 2018). bvFTD affects not only regional power but the organisation of resting activity. Greater low-frequency burden may reflect reduced cortical network efficiency or impaired maintenance of fast resting states. Reduced beta residual power indicates that the loss of beta is not simply a consequence of a changing 1/f background but includes reduced beta structure after aperiodic correction. Increased within-source alpha-phase/beta-amplitude PAC, including in LAI and ACC, suggests altered cross-frequency organisation in regions involved in salience processing, behavioural control, and switching between internal and external modes. This does not prove a specific cellular mechanism, but it provides a physiological phenotype of bvFTD. While such metrics may provide biomarkers for treatment outcomes (Hughes et al., 2025), they do not explain the causes of abnormal physiology and behavior. Phase-amplitude coupling (alpha-phase/beta-amplitude) quantified the dependence of beta amplitude on alpha phase within each source (Canolty et al., 2006; Tort et al., 2010). The increase in bvFTD is consistent with altered cross-frequency organisation, not an absence of alpha-phase/beta-amplitude coupling; its functional significance and any role in hierarchical control remain hypotheses rather than established mechanisms. Despite significant group-level PAC differences, classification based on panels containing PAC and other spectral features remained modest in this exploratory analysis.

Previous studies have indicated partial overlap in the clinical, spectral and topographical changes of PSP and bvFTD (Sami et al., 2018; Ghosh et al., 2012; Murley et al., 2020). PSP clinical data were coherent at the individual level (accuracy >0.93), but the spectral abnormalities were weakly coherent (accuracy <0.6). This indicates that PSP-related resting-state MEG changes are distributed, lower amplitude, or expressed in subcortical systems not fully represented by our cortical sources. In contrast, the DCM classification provided a stronger signal, with a distributed latent pattern of abnormal timing, coupling and gain.

The comparison of bvFTD-vs-PSP highlights the importance of aperiodic features. The 1/f slope survived FDR correction across all seven sources in classifying patients. A steeper 1/f slope can be interpreted as a field-potential marker of altered asynchronous population activity or excitation-inhibition balance (Gao et al., 2017; Donoghue et al., 2020). However, the individual-level PSD classification between diseases was modest (accuracy <0.640). DCM panel that distinguished patients was compact, including ACC-to-MPFC NMDA coupling, MPFC-to-LAI NMDA feedback, and extrinsic conduction delays. This suggests that bvFTD and PSP differ in salience/default-mode timing and receptor-specific connectivity, without singular receptor abnormalities.

Pooling over patient groups may identify shared disease burden. However, for bvFTD and PSP, pooling weakened the physiological specificity and greatly reduced the accuracy of physiological classification despite high clinical classification. In other words, similar cognitive deficits arise from distinct pathophysiological abnormalities between the disorders.

The comparison between linear and nonlinear classifiers helps to interpret the geometry of diseases. Linear SVM performance in several DCM contrasts suggests that disease differences can be expressed as weighted combinations of weak features. The occasional advantage of random forest and AdaBoost classifiers in PSD-based contrasts, raises the possibility of feature interactions or threshold effects. However, flexible classifiers are more vulnerable to overfitting in modest samples, particularly when MCC and AUC are low.

This study has limitations. The group-to-individual assay examines how representations identified by group-level analyses support individual assignment within the study cohort. Classification functions serve as probes of this correspondence. Feature panels were constructed from the full-sample group statistics and held fixed while each participant was excluded from fitting the decision function. So as a first limitation, the reported metrics may be optimistically biased, and describing the analysis as a coherence assay rather than a diagnostic test. Independent evaluation of individual-level generalisation would require feature selection, scaling, hyperparameter optimisation, model selection, and threshold selection to be nested fully within the training data of each validation fold, followed by independent external validation. Second, the cohort size is large for MEG studies, but modest for machine learning purposes. Although some features differed by large (>1) or very large (>2) effect sizes, this does not remove selection bias or establish out-of-sample generalisability. Third, classification performance might be increased by post-feature-selection tuning (e.g., sequential forward selection or embedding techniques), hyperparameter optimisation, alternative decision thresholds, probability calibration, and model-specific threshold selection. Fourth, our study was from a single site. Larger and longitudinal studies with external validation will help to determine whether the spectral and DCM signatures generalise across sites, scanners, disease stages, and progression trajectories. Note too, that clinical scores and MEG features can answer different questions: strong clinical performance does not invalidate MEG, but the focus of MEG remains currently on physiological interpretation and mechanistic value rather than diagnosis.

The translation of group-to-individual features reduces the risk of two opposing errors as it avoids treating group differences as sufficient for an individual’s biomarker, and it avoids overlooking physiologically informative features that are diverse across individuals. Clinical scales indicate how strongly the syndrome is expressed; PSD indicates the alteration of resting spectral organisation; and biophysical modelling helps to interpret those alterations in terms of latent physiological mechanisms. The clinical, spectral, and mechanistic levels of description are complementary for individualised phenotyping. The trade-off between mechanistic insight and classification can improve the design of much-needed experimental medicine and clinical trials, for people with neurodegenerative disorders.

## Data and Methods

### Participants and clinical phenotyping

We analysed resting-state MEG recordings from three groups: healthy controls (CON; N=74), people with bvFTD (N=45), and people with PSP (N=55), and the pooled patient group (bvFTD+PSP). Demographic and clinical data are summarised in Table 1. Diagnoses were made according to contemporary clinical criteria for probable bvFTD (Rascovsky et al., 2011) and PSP (Höglinger et al., 2017; Richardson’s syndrome), supported by structural MRI. Clinical evaluation included Addenbrooke’s Cognitive Examination-Revised (ACE-R), Mini-Mental State Examination (MMSE), Frontotemporal Dementia Rating Scale (FRS), Progressive Supranuclear Palsy Rating Scale (PSPRS), Frontal Assessment Battery (FAB), and Cambridge Behavioural Inventory-Revised (CBI-R). These were analysed across the four contrasts (CON vs bvFTD, CON vs PSP, bvFTD vs PSP, and CON vs bvFTD+PSP). Clinical scores provide a benchmark for features that are expected to be individually coherent because they include the defining behavioural, cognitive, and motor deficits of each disease.

### Imaging and MEG

Participants underwent 5 minutes’ resting-state MEG recording with eyes open using an Elekta VectorView system with 204 planar gradiometers and 102 magnetometers, sampled at 1000 Hz. Eye movements were recorded using bipolar electrooculogram. Head position was tracked using five head-localisation indicator coils mounted on the EEG cap, and Polhemus digitisation recorded three fiducial points together with more than 100 head-shape points. Preprocessing used the Elekta Neuromag toolbox, MaxFilter v2.2.1. Bad sensors were identified and interpolated, temporal signal space separation was applied for noise reduction, and head movement was corrected. Data was downsampled to 500 Hz, band-pass filtered from 0.1 to 100 Hz, notch filtered at 48-52 Hz and 98-102 Hz, cleaned with ICA informed by EOG channels, segmented into 1 s epochs, and subjected to epoch-wise artefact rejection.

For each participant, a T1-weighted structural MRI scan was acquired on a 3 T Siemens system using an MPRAGE sequence (TR=2300 ms, TE=2.91 ms, 1 mm isotropic resolution) or a 7T structural was acquired on a Siemens TERRA system (Siemens Healthineers, Erlangen, Germany) with 32-channel head coil. Acquisition parameters were: 0.75 mm isotropic voxels, echo time (TE) = 1.99 ms, repetition time (TR) = 4300 ms, inversion times = 840 ms/2370 ms) and converted from DICOM to NIfTI format. Source inversion used SPM12’s Bayesian framework with a single-shell forward model constructed from participant-specific anatomy. The inversion was constrained to 0.1-100 Hz across all epochs and fused magnetometer and gradiometer data during estimation. Source time courses were then extracted from the seven SN-DMN regions: bilateral anterior insula (LLAI, RAI), anterior cingulate cortex (ACC), medial prefrontal cortex (MPFC), posterior cingulate cortex (PCC), bilateral angular gyrus (LAG, RAG).

Power spectral density was estimated for each source and epoch using Welch’s method. Log-transformed power spectra were used for visualisation and frequency-wise group comparison. Features were chosen to represent interpretable aspects of resting physiology such as low-frequency burden, alpha and beta organisation, aperiodic 1/f structure, spectral dispersion, peak frequency, and alpha-phase/beta-amplitude coupling. The 1/f slope was estimated in log-log spectral space and treated as an indirect field-potential marker of aperiodic spectral steepness, asynchronous population activity, and possible excitation-inhibition imbalance. Spectral entropy and median frequency summarised spectral dispersion and cumulative frequency organisation, with lower values supporting a slower or less diverse spectral state when combined with other features. Alpha-phase/beta-amplitude coupling (PAC) quantified the dependence of beta amplitude on alpha phase within each source and was interpreted as an index of within-source cross-frequency organisation, not between-region connectivity or directed influence. Higher PAC indicates stronger phase-dependent variation in beta amplitude but does not by itself establish more effective coordination or a specific cellular mechanism.

### Dynamic causal modelling

We fitted a conductance-based Dynamic Causal Model (aka CMM-NMDA) to the seven SN-DMN sources using the cross-spectral-density DCM framework implemented in SPM12 (Friston et al., 2003; Marreiros et al., 2009; Litvak et al., 2011; Moran et al., 2013). Participant-specific model inversion from MEG data, used the variational Bayesian scheme, yielding posterior means for intrinsic microcircuit couplings, extrinsic between-source AMPA/NMDA coupling terms, membrane and synaptic time constants, and conduction delays (Appendix A).

We use descriptive labels for the CMM-NMDA parameter families. Regions are LAI, ACC, RAI, MPFC, PCC, LAG, and RAG. Population labels are SE (spiny excitatory), SP (superficial pyramidal), IN (inhibitory), and DP (deep pyramidal) and explained in Appendix A. Between-region AMPA coupling is written 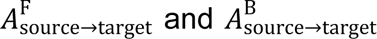 for forward and backward connections, respectively; 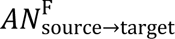 and 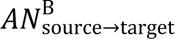 denote the corresponding NMDA couplings. F/B specifies the modelled connection class, while the arrow specifies anatomical direction. Within-region coupling is written 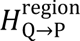, from population Q to population P in the named region. Synaptic time constants are written 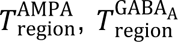, and 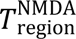 for AMPA-mediated excitatory, GABA_A_-mediated inhibitory, and NMDA-mediated excitatory time constants. *CV*_population_ denotes membrane capacitance of the named population; for example, *CV*_SP_ refers to superficial pyramidal cells. *D*_int_ and *D*_ext_ denote intrinsic and extrinsic delays, respectively. The terms 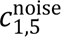 and 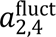 denote the specific sensor-noise and endogenous-fluctuation coefficients.

### Statistical analysis and group-to-individual translation assay

All analyses were performed for four contrasts: controls-bvFTD+PSP, controls-bvFTD, controls-PSP, and bvFTD-PSP. Univariate group comparisons used Welch’s two-sample t-test to allow for unequal variances (Welch, 1947). For each clinical score, PSD feature, and inferred DCM parameter, we report p-values together with standardised effect sizes, including pooled-standard-deviation Cohen’s d (Cohen, 1988) and bias-corrected Hedges’ g (Hedges, 1981). Multiple comparisons were controlled within each analysis family using Benjamini-Hochberg false-discovery-rate correction (Benjamini and Hochberg, 1995).

Because the feature spaces were expected to be high-dimensional and redundant, we developed feature-ranking panels using full-sample group statistics before the group-to-individual tests (Table 2) commonly referred as filter rankings (Guyon and Elisseeff, 2003). Four ranking approaches were used at panel sizes of five and ten features: P5/P10 selected the smallest uncorrected p-values, D5/D10 selected the largest absolute Cohen’s d values, C5/C10 combined statistical evidence and effect size, and S5/S10 added a recurrence bonus across contrasts. C and S are study-defined composite scores. These panels are group-derived summaries of statistically prominent effects used to probe their collective contribution to individual assignment. Statistical significance alone does not determine predictive value (Lo et al., 2015).

**Table 2.** Summary of feature-ranking panels used in the group-to-individual translation assay.

| Panel | Selection rule | What it emphasizes | Strength | Limitation |
| --- | --- | --- | --- | --- |
| P5 | 5 smallest uncorrected p-values | Strongest univariate statistical evidence | Compact and transparent | May prioritise small but precisely estimated effects |
| P10 | 10 smallest uncorrected p-values | Broader statistically ranked set | Wider coverage of candidate effects | Greater risk of adding weak or noisy features |
| D5 | 5 largest absolute Cohen's d values | Strongest standardised marginal group separation | Emphasises effect magnitude | Effect sizes can be unstable in modest samples |
| D10 | 10 largest absolute Cohen's d values | Broader large-effect panel | Broader coverage of large marginal effects | More vulnerable to noisy large-effect terms |
| C5 | 5 largest C scores (Equation 2) | Equal weighting after cross-feature standardisation | Integrates two ranking criteria | Study-defined weights; does not assess redundancy |
| C10 | 10 largest C scores (Equation 2) | Broader combined-score panel | Integrates two ranking criteria | Can add weak or redundant features |
| S5 | 5 largest S scores (Equation 3) | Combined score plus $0.50 \times$ normalised recurrence | Favours recurring group-ranked features | Depends on contributing contrasts; shared participants |
| S10 | 10 largest S scores (Equation 3) | Broader panel with the same recurrence bonus | Retains a wider set of recurring features | May mix contrast-specific and shared disease effects |

For feature j in contrast c, let p denote its uncorrected Welch p-value and d its pooled-SD Cohen’s d. The P and D scores were defined as

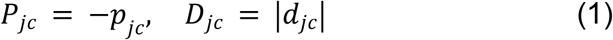

The combined score adds equally weighted, standardised evidence and effect-size components:

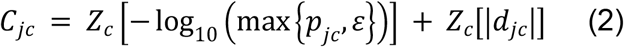

Here Z is a z-score calculated across eligible matched features within the same feature domain and contrast, using the sample mean and sample standard deviation. This standardises the ranking statistics across features, separately from participant-level predictor scaling

For recurrence, R counts the number of contributing contrasts in which a feature appears among the first 20 matching entries in the relevant ranked list. The count is normalised by the largest observed recurrence across candidate features, and the S score is

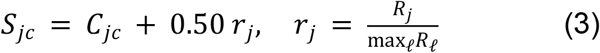

We used leave-one-out classification as a probe of the individual-level expression of group-defined feature panels. The aim was to examine how features prioritised by group-level statistics, when considered together, support assignment of individual participants within the study cohort. By design, feature panels were defined using the full-sample group statistics and held fixed across classification folds. In each fold, the left-out participant was excluded from classifier fitting. This preserves the representation identified by the group analysis while examining individual assignments made by decision functions fitted without that participant.

Classifier performance was evaluated using accuracy, sensitivity, specificity, precision, F1-score, Matthews correlation coefficient (MCC) and area under the receiver operating characteristic curve (AUC). These metrics serve as descriptors of the features that indicate, for example, whether a group effect is distributed consistently across participants or concentrated in a subset, and whether discriminative information is balanced across classes (sensitivity versus specificity) or carried asymmetrically. Because panel definition and classification share the same sample, these metrics characterise the selected panels conditional on the group analysis and are not intended as unbiased estimates of out-of-sample generalisation or prospective diagnostic performance (Ambroise and McLachlan, 2002).

We evaluated linear support vector machine (linear SVM), radial-basis-function support vector machine (RBF-SVM), random forest, and AdaBoost in this study. These models were used as analytic probes of how group-derived feature panels are formed at the individual level. All classifiers were applied to the same group-derived feature panels within each contrast so that differences in performance could be interpreted in relation to the geometry of the selected feature space.

The linear SVM tests whether the selected features separate groups along an approximately additive, weighted axis. Within the full-sample-selected representation, strong linear-SVM performance is consistent with individual-level separation by a relatively simple combination of feature values. The RBF-SVM allows curved decision boundaries and is sensitive to nonlinear separation, making it useful for detecting feature-space structure that cannot be represented by a single linear axis. Random forest and AdaBoost are tree-based ensemble methods. Random forest captures interactions and variable-dependent thresholds through averaging across multiple decision trees, whereas AdaBoost sequentially emphasises observations and can reveal whether a small set of decision rules improves assignment. The exploratory comparison across classifier families examines whether (i) compact panels perform well with a linear model, or (ii) the signal is relatively sparse, non-linear and/or additive. The comparison of classifier families also provides a formal interpretation of the group-to-individual translation assays.

The analysis can be represented as a composition of mappings where group-level contrasts define candidate feature panels; each participant is represented by a feature vector within a selected panel; and the classifier maps that vector to a decision score and binary assignment evaluated using leave-one-out classification. In category-theoretic terms, these composable mappings are morphisms linking population-level feature characterisation to individual-level assignment. This terminology describes the mathematical workflow where a morphism denotes a composable mapping. For the linear SVM, the decision function is affine-linear, such that assignment is determined by an additive weighted combination of the selected features. Strong performance of this model would therefore be consistent with an approximately linear group-separating axis in the selected representation. By contrast, the RBF-SVM produces a decision function that is nonlinear in the original feature space, whereas random forest and AdaBoost implement piecewise mappings based on feature interactions and thresholds. Better leave-one-out performance from these nonlinear machine learning tools would motivate hypotheses about curved decision boundaries, interactions, or heterogeneous feature configurations within the selected representation.

## Data availability and Code availability

Enquiries about access to the underlying participant data should be directed to the senior author, Professor James B. Rowe. The analysis code used in this study will be made publicly available upon publication.

## Acknowledgements

This work is primarily funded by the Wellcome Trust (220258) with additional support from Dementias Platform UK (funded by the Medical Research Council MC_UU_00030/14 & MR/T033371/1, Janssen, Eli Lilly, AstraZeneca, Araclon, IXICO, Somalogic, GlaxoSmithKline, Invicro, Cambridge Cognition, and Cognetivity), Alzheimers Research UK (ARUK-PG2017B-19), NIHR Cambridge Biomedical Research Centre (NIHR203312) and National Institute for Health Research Oxford Health Biomedical Research Centre (NIHR203316). The views expressed are those of the authors and not necessarily those of the NIHR or the Department of Health and Social Care. For the purpose of open access, the authors have applied a CC. BY public copyright licence to any Author Accepted Manuscript version arising from this submission.

## Author Contributions

**AJ**: conceptualization, methodology, software, validation, formal analysis, data curation, finalizing research program and writing original drafts, **LH**: Study design, MEG data collection, clinical scores / cognitive assessments, **MN, MAR**: data collection and reading final draft of paper upon completion of research, **SH**: reading final draft of paper upon completion of research, **TC, NH, AM, TR, DJW**: clinical diagnostics, reading final draft of paper upon completion of research; **JR**: discussion, methodology, original draft, funding acquisition.

