## appendices A to D for "From group-level neurophysiological biomarkers to individual-level phenotyping to identify mechanisms of neurodegenerative disease"

#### Appendix A. CMM-NMDA Model Parameters and DCM Statistical Results

This appendix defines the source-region indices, neuronal-population indices, and parameter-family notation used for the CMM-NMDA dynamic causal modelling analyses.

##### Region index

| Index | Region |
|---|---|
| 1 | Left anterior insula (LAI) |
| 2 | ACC |
| 3 | Right anterior insula (RAI) |
| 4 | MPFC |
| 5 | PCC |
| 6 | LAG |
| 7 | RAG |

##### Population index

| Index | Population |
|---|---|
| 1 | Spiny excitatory |
| 2 | Superficial pyramidal |
| 3 | Inhibitory |
| 4 | Deep pyramidal |

##### Parameter-family definitions

| Parameter family | Definition |
|---|---|
| D(1), D(2) | intrinsic delay; extrinsic delay. |
| S(1) | NMDA-mediated population variance. |
| T(r,k) | time constants in region r with k = 1 intrinsic excitatory AMPA, k = 2 inhibitory GABA_A, and k = 3 excitatory NMDA. |
| CV(c) | membrane capacitance parameter for population c, where c in {1, 2, 3, 4}. |
| E(1) | background activity. |
| A{1}(i,j) | forward AMPA extrinsic coupling from region j to region i. |
| A{2}(i,j) | backward AMPA extrinsic coupling from region j to region i. |
| AN{1}(i,j) | forward NMDA extrinsic coupling from region j to region i. |
| AN{2}(i,j) | backward NMDA extrinsic coupling from region j to region i. |
| H(p,q,n) | intrinsic microcircuit coupling within region n from population q to population p, where p, q in {1, 2, 3, 4} and n in {1, ..., 7} indexes regions. |
| L(n) | electrode gain for region n. |
| J(3) and J(4) | contributions of spiny and deep layers to LFP, respectively. |
| *a* | Endogenous random fluctuation with transfer function $a_1/\omega^2$. |
| *d* | Structural cosine coefficients of endogenous random fluctuation. |
| *b* | Common sensor noise with transfer function $b_1/\omega^2$. |
| *c* | Specific sensor noise with transfer function $c_1/\omega^2$. |
| *f* | Scaling some frequencies as model of data filtration. |

##### A.2. Statistical test for DCM parameter comparisons

This section adds the DCM-parameter statistical results. Each candidate parameter was tested with Welch's two-sample t-test, standardised effect sizes were reported as Cohen's d and Hedges' g, and Benjamini-Hochberg false-discovery-rate correction. Across all four DCM contrasts, with 337 inferred parameters, no DCM parameter survived BH-FDR correction at $q < 0.05$. Consequently, the tables below should be read as top-ranked uncorrected or trend-level effects.

**Table A1. DCM univariate statistical test summary across contrasts. p-values are uncorrected Welch-test p-values; q-values are BH-FDR adjusted within the DCM parameter family.**

| Contrast | N1 / N2 | Valid tests | p<0.05 | p<0.01 | q<0.05 | min p | min q |
|---|---|---|---|---|---|---|---|
| CON vs bvFTD | 74 / 45 | 337 | 16 | 4 | 0 | 8.35e-04 | 0.281 |
| CON vs PSP | 74 / 55 | 337 | 21 | 1 | 0 | 0.00685 | 0.667 |
| CON vs pooled patients | 74 / 100 | 337 | 24 | 6 | 0 | 0.00282 | 0.410 |
| bvFTD vs PSP | 45 / 55 | 337 | 15 | 3 | 0 | 0.002 | 0.723 |

**Table A2. Dominant top-ranked DCM parameter families and contrast-level interpretation. Counts refer to the ten lowest uncorrected p-values in each contrast.**

| Contrast | Dominant top-10 families | Interpretation |
|---|---|---|
| CON vs bvFTD | intrinsic microcircuit coupling (4); forward AMPA coupling (2); delay (1); synaptic time constant (1); membrane capacitance (1); backward AMPA coupling (1) | Uncorrected pattern concentrated in extrinsic delay, RAG AMPA timing, superficial-pyramidal capacitance/self-coupling, and AMPA communication. |
| CON vs PSP | intrinsic microcircuit coupling (3); backward NMDA coupling (2); forward AMPA coupling (1); specific sensor noise (1); endogenous fluctuation (1); membrane capacitance (1); forward NMDA coupling (1) | Uncorrected effect is weaker than the bvFTD-control contrast and are distributed across PCC/LAI/MPFC intrinsic coupling, AMPA links, NMDA terms, capacitance, and modelled fluctuation/noise terms. |
| CON vs pooled patients | forward AMPA coupling (4); intrinsic microcircuit coupling (3); membrane capacitance (1); synaptic time constant (1); specific sensor noise (1) | Patient pooled contrast shows the largest number of uncorrected effects, dominated by superficial-pyramidal capacitance/self-coupling and forward AMPA communication. |
| bvFTD vs PSP | backward NMDA coupling (3); structural cosine coefficient (2); synaptic time constant (1); forward NMDA coupling (1); forward AMPA coupling (1); delay (1); electrode gain (1) | Direct syndrome pattern emphasises RAG excitatory timing, ACC-to-MPFC NMDA coupling, LAG-to-RAG AMPA coupling, extrinsic delay, MPFC-to-LAI NMDA feedback, and posterior gain/fluctuation terms. |

**Table A3. DCM group-to-individual classification results.**

| Contrast | Best reported DCM model/panel | Headline performance | Physiological interpretation |
|---|---|---|---|
| CON vs bvFTD | Linear SVM, ten-parameter panel | Accuracy 0.748; specificity 0.878; MCC 0.446; AUC 0.756 | Moderate individual-level coherence; distributed timing, capacitance/local-gain, and AMPA-mediated SN-DMN communication rather than a single-parameter effect. |
| CON vs PSP | Linear SVM, P10 panel | Accuracy 0.698; sensitivity 0.491; specificity 0.851; precision 0.711; F1 0.581; MCC 0.371; AUC 0.770 | Moderate DCM coherence, stronger than the CON-PSP PSD classifier result; interpreted as a distributed latent timing/gain/coupling pattern rather than a corrected single-parameter biomarker. |
| CON vs pooled patients | Random Forest P10/STABLE_TOP10 by accuracy/F1/MCC; linear SVM compact five-parameter panel by AUC | RF: accuracy 0.632; sensitivity 0.720; specificity 0.514; F1 0.692; MCC 0.238; AUC 0.639. Linear-SVM compact panel: AUC 0.648; accuracy 0.598; MCC 0.174 | Weak-to-modest common disease-control axis; pooling preserves some shared gain/coupling effects but dilutes syndrome-specific physiology. |
| bvFTD vs PSP | Linear SVM, compact five-parameter DCM panel | Accuracy 0.740; PSP sensitivity 0.800; bvFTD specificity 0.667; precision 0.746; F1 0.772; MCC 0.472; AUC 0.798 | Moderate syndrome-level coherence; compact timing/coupling panel involving RAG excitatory timing, ACC-MPFC NMDA coupling, LAG-RAG AMPA coupling, MPFC-LAI NMDA feedback, and extrinsic delay. |

Selected parameters in the reported DCM classification panels:

- CON vs bvFTD: D(2), T(7,1), H(1,1,2), CV(2), H(2,2,2), A{1}(3,7), H(2,2,3), A{1}(5,2), A{2}(1,3), H(2,2,4)
- CON vs PSP: H(1,3,5), A{1}(4,3), H(2,2,4), c(1,5), a(2,4), H(2,3,1), CV(2), AN{2}(1,4), AN{2}(4,6), AN{1}(7,2)
- CON vs pooled patients: P10/STABLE_TOP10: CV(2), H(2,2,4), A{1}(4,3), H(2,2,2), A{1}(5,2), T(1,2), H(1,1,2), A{1}(4,5), c(1,5), A{1}(3,7). Compact five-parameter AUC panel: CV(2), H(2,2,4), A{1}(4,3), H(2,2,2), A{1}(5,2)
- bvFTD vs PSP: T(7,1), AN{1}(4,2), A{1}(7,6), AN{2}(1,4), D(2)

##### A.2.1. CON vs bvFTD

Controls-bvFTD showed the strongest DCM-supported patient-control pattern. Positive Cohen's d values indicate higher posterior estimates in controls, whereas negative values indicate higher estimates in bvFTD. The pattern combines extrinsic transmission delay, RAG AMPA-mediated time constants, superficial-pyramidal capacitance, ACC/RAI/MPFC intrinsic self-coupling, and AMPA-mediated communication across posterior, cingulate, and bilateral anterior insular nodes. Because no single parameter survives FDR correction, the result is best interpreted as a distributed physiological signature rather than a decisive isolated abnormality.

**Table A4. CON-bvFTD: top ten uncorrected DCM parameter effects.**

| Rank | Parameter | Higher mean | Mean A | Mean B | p_Welch | q_BH | Cohen's d |
|---|---|---|---|---|---|---|---|
| 1 | D(2) | CON | 0.234 | 0.027 | 8.35e-04 | 0.281 | 0.64 |
| 2 | T(7,1) | CON | 0.038 | -0.027 | 0.003 | 0.520 | 0.51 |
| 3 | CV(2) | bvFTD | 0.152 | 0.373 | 0.006 | 0.657 | -0.55 |
| 4 | H(1,1,2) | CON | -0.008 | -0.157 | 0.008 | 0.657 | 0.60 |
| 5 | H(2,2,2) | CON | 0.126 | -0.038 | 0.012 | 0.730 | 0.57 |
| 6 | A{1}(3,7) | bvFTD | 0.011 | 0.205 | 0.015 | 0.730 | -0.54 |
| 7 | H(2,2,3) | CON | 0.014 | -0.190 | 0.015 | 0.730 | 0.51 |
| 8 | A{2}(1,3) | CON | 0.052 | -0.046 | 0.020 | 0.787 | 0.44 |
| 9 | A{1}(5,2) | bvFTD | -0.004 | 0.096 | 0.021 | 0.787 | -0.51 |
| 10 | H(2,2,4) | CON | 0.047 | -0.105 | 0.026 | 0.839 | 0.46 |

##### A.2.2. CON vs PSP

Controls-PSP showed weaker univariate DCM effects than the bvFTD-control contrast. One parameter reached uncorrected $p < 0.01$, 21 parameters reached uncorrected $p < 0.05$, and none survived FDR correction. The leading parameters include PCC inhibitory-to-spiny coupling, RAI-to-MPFC AMPA coupling, MPFC superficial-pyramidal self-coupling, a specific sensor-noise term, endogenous fluctuation, LAI inhibitory-to-superficial-pyramidal coupling, membrane capacitance, and NMDA-mediated long-range coupling terms. These effects should be treated as distributed trend-level physiology rather than corrected single-parameter biomarkers. The best CON-PSP result was a linear SVM using the P10 panel, with accuracy 0.698, sensitivity 0.491, specificity 0.851, precision 0.711, F1-score 0.581, MCC 0.371, and AUC 0.770.

**Table A5. CON vs PSP: top ten uncorrected DCM parameter effects.**

| Rank | Parameter | Higher mean | Mean A | Mean B | p_Welch | q_BH | Cohen's d |
|---|---|---|---|---|---|---|---|
| 1 | H(1,3,5) | CON | 0.080 | -0.009 | 0.00685 | 0.667 | 0.486 |
| 2 | A{1}(4,3) | PSP | 0.136 | 0.409 | 0.016 | 0.667 | -0.465 |
| 3 | H(2,2,4) | CON | 0.047 | -0.122 | 0.018 | 0.667 | 0.460 |
| 4 | c(1,5) | CON | -0.030 | -0.048 | 0.0203 | 0.667 | 0.443 |
| 5 | a(2,4) | PSP | 0.237 | 0.424 | 0.0224 | 0.667 | -0.440 |
| 6 | H(2,3,1) | CON | 0.032 | 0.000 | 0.0284 | 0.667 | 0.380 |
| 7 | CV(2) | PSP | 0.152 | 0.327 | 0.0308 | 0.667 | -0.404 |
| 8 | AN{2}(1,4) | CON | 0.053 | -0.017 | 0.0325 | 0.667 | 0.364 |
| 9 | AN{2}(4,6) | CON | 0.022 | -0.015 | 0.0331 | 0.667 | 0.352 |
| 10 | AN{1}(7,2) | PSP | -0.020 | 0.035 | 0.0353 | 0.667 | -0.417 |

##### A.2.3. CON vs pooled patients

The pooled patient-control comparison used 74 controls and 100 pooled patients. It contained the largest number of uncorrected DCM effects among the contrasts, but the minimum FDR-adjusted q-value remained far above 0.05. The leading parameters were recurrent across syndrome-specific contrasts, especially superficial-pyramidal capacitance and self-coupling, forward AMPA links involving RAI, MPFC, PCC, ACC, and RAG, LAI inhibitory timing, ACC intrinsic excitatory coupling, and a specific sensor-noise component. The pooled-patient individual-level coherence remained weak-to-modest where the strongest operating point by accuracy, F1-score, and MCC was a Random Forest using the P10/STABLE_TOP10 panel (accuracy 0.632; sensitivity 0.720; specificity 0.514; F1-score 0.692; MCC 0.238; AUC 0.639), while the highest AUC was obtained by a linear SVM using the compact five-parameter panel [CV(2), H(2,2,4), A{1}(4,3), H(2,2,2), A{1}(5,2)] (AUC 0.648; accuracy 0.598; MCC 0.174).

**Table A6. CON vs pooled patients: top ten uncorrected DCM parameter effects.**

| Rank | Parameter | Higher mean | Mean A | Mean B | p_Welch | q_BH | Cohen's d |
|---|---|---|---|---|---|---|---|
| 1 | CV(2) | bvFTD,PSP | 0.152 | 0.348 | 0.00282 | 0.410 | -0.452 |
| 2 | H(2,2,4) | CON | 0.047 | -0.114 | 0.003 | 0.410 | 0.432 |
| 3 | A{1}(4,3) | bvFTD,PSP | 0.136 | 0.388 | 0.005 | 0.410 | -0.406 |
| 4 | H(2,2,2) | CON | 0.126 | 0.015 | 0.006 | 0.410 | 0.405 |
| 5 | A{1}(5,2) | bvFTD,PSP | -0.004 | 0.070 | 0.006 | 0.410 | -0.403 |
| 6 | A{1}(4,5) | bvFTD,PSP | -0.068 | 0.046 | 0.009 | 0.497 | -0.371 |
| 7 | H(1,1,2) | CON | -0.008 | -0.098 | 0.012 | 0.535 | 0.367 |
| 8 | A{1}(3,7) | bvFTD,PSP | 0.011 | 0.150 | 0.016 | 0.535 | -0.346 |
| 9 | T(1,2) | bvFTD,PSP | 0.078 | 0.146 | 0.016 | 0.535 | -0.357 |
| 10 | c(1,5) | CON | -0.030 | -0.044 | 0.017 | 0.535 | 0.353 |

##### A.2.4. bvFTD vs PSP

The direct syndrome comparison used 45 bvFTD and 55 PSP participants. No DCM parameter survived BH-FDR correction, but the uncorrected pattern was compact and physiologically interpretable. The leading effects were centered on RAG AMPA-mediated excitatory timing [T(7,1)], forward NMDA coupling from ACC to MPFC [AN{1}(4,2)], forward AMPA coupling from LAG to RAG [A{1}(7,6)], extrinsic delay [D(2)], and backward NMDA coupling from MPFC to LAI [AN{2}(1,4)]. PSP showed higher mean values for T(7,1), A{1}(7,6), and D(2), whereas bvFTD showed higher mean values for AN{1}(4,2) and AN{2}(1,4). The top ten also included structural cosine coefficients of endogenous fluctuations [d(2,7), d(3,7)], posterior gain [L(5)], and additional backward NMDA terms; these should be interpreted cautiously because they include modelled fluctuation/observation components as well as synaptic coupling terms.

A linear SVM using the compact five-parameter panel [T(7,1), AN{1}(4,2), A{1}(7,6), AN{2}(1,4), D(2)] achieved accuracy 0.740, PSP sensitivity 0.800, bvFTD specificity 0.667, precision 0.746, F1-score 0.772, MCC 0.472, and AUC 0.798. This indicates moderate individual-level coherence in DCM space for the direct syndrome contrast.

**Table A7. bvFTD vs PSP: top ten uncorrected DCM parameter effects.**

| Rank | Parameter | Higher mean | Mean A | Mean B | p_Welch | q_BH | Cohen's d |
|---|---|---|---|---|---|---|---|
| 1 | T(7,1) | PSP | -0.027 | 0.036 | 0.002 | 0.723 | -0.622 |
| 2 | AN{1}(4,2) | bvFTD | 0.148 | -0.055 | 0.005 | 0.781 | 0.634 |
| 3 | A{1}(7,6) | PSP | -0.031 | 0.166 | 0.009 | 0.797 | -0.526 |
| 4 | D(2) | PSP | 0.027 | 0.232 | 0.012 | 0.797 | -0.494 |
| 5 | AN{2}(1,4) | bvFTD | 0.143 | -0.017 | 0.013 | 0.797 | 0.556 |
| 6 | d(2,7) | bvFTD | 0.008 | -0.000 | 0.020 | 0.797 | 0.488 |
| 7 | AN{2}(5,7) | PSP | -0.006 | 0.012 | 0.021 | 0.797 | -0.446 |
| 8 | L(5) | bvFTD | 0.294 | 0.179 | 0.021 | 0.797 | 0.492 |
| 9 | AN{2}(7,6) | PSP | -0.063 | 0.041 | 0.023 | 0.797 | -0.454 |

| Rank | Parameter | Higher mean | Mean A | Mean B | p_Welch | q_BH | Cohen's d |
|---|---|---|---|---|---|---|---|
| 10 | d(3,7) | bvFTD | 0.007 | -0.002 | 0.029 | 0.797 | 0.452 |

### Appendix B. Statistical test results for PSD feature comparisons

#### B.1. PSD feature comparisons

This appendix explains the feature-level statistical-test output. The tables below summarise the strongest univariate PSD effects for each contrast. Each contrast contains 105 PSD-derived features tested with Welch's two-sample t-test. We report group means, standard deviations, uncorrected p-values, Benjamini-Hochberg false-discovery-rate adjusted q-values, and Cohen's d. Effects with q_BH < 0.05 are treated as corrected feature-level effects; effects with p_Welch < 0.05 but q_BH >= 0.05 are reported as uncorrected or trend-level effects and should be interpreted cautiously.

#### B.2. Channel and features information

**Table B1. Source-channel mapping used in the PSD statistical analysis.**

| Workbook channel | Source region | Network interpretation |
|---|---|---|
| Ch1 | Left anterior insula (LAI) | Salience-network source |
| Ch2 | ACC | Salience-network source |
| Ch3 | Right anterior insula (RAI) | Salience-network source |
| Ch4 | MPFC | Default-mode/posterior association source |
| Ch5 | PCC | Default-mode/posterior association source |
| Ch6 | LAG | Default-mode/posterior association source |
| Ch7 | RAG | Default-mode/posterior association source |

**Table B2. PSD feature families and interpretation.**

| Feature family | Operational meaning | Physiological interpretation in this paper |
|---|---|---|
| Relative delta/theta | Proportion of total power in slow bands. | Higher values suggest greater low-frequency burden or slowing. |
| Relative alpha/beta/gamma | Band-limited power normalised by total power. | Lower beta or gamma can indicate weaker organised fast rhythmic activity. |
| Alpha/beta residual max | Peak oscillatory residual after accounting for aperiodic background. | Lower beta residual supports reduced beta rhythmic structure beyond broad 1/f tilt. |
| 1/f slope | Aperiodic spectral slope in log-log power-frequency space. | More negative slope indicates broader aperiodic redistribution; interpretation should remain cautious. |
| Spectral entropy/median frequency | Summary of spectral dispersion and cumulative frequency distribution. | Lower values generally indicate less diverse or slower spectral organisation. |
| PAC alpha phase-beta amplitude | Modulation of beta amplitude by alpha phase. | Altered PAC suggests disturbed cross-frequency coordination, especially in SN sources. |
| Band ratios | Low-frequency power divided by alpha power. | Higher ratios suggest slowing relative to organised alpha activity. |

In the contrast tables, “higher mean” identifies the group with the larger numeric mean. For the 1/f slope a larger value is less negative, whereas the manuscript interpretation focuses on bvFTD showing a more negative slope relative to PSP.

#### B.3. Summary across contrasts

**Table B3. Statistical-test summary across the four PSD feature contrasts.**

| Contrast | No. features | p<0.05 | p<0.01 | q<0.05 | min p | min q | Dominant families | Interpretation |
|---|---|---|---|---|---|---|---|---|
| CON vs bvFTD | 105 | 37 | 20 | 20 | 7.39e-04 | 0.029 | PAC alpha-phase/beta-amp (7 corrected); Beta residual max (7 corrected); Relative beta (3 corrected); Spectral entropy (3 corrected) | Corrected PSD signature: higher PAC and lower beta residual/relative beta/spectral entropy in bvFTD. |
| CON vs PSP | 105 | 1 | 0 | 0 | 0.041 | 0.763 | 1/f slope (5 top-ranked); Beta peak frequency (5 top-ranked) | No corrected univariate PSD effect; only weak aperiodic/beta-peak trends, with weak individual-level PSD classification. |
| bvFTD vs PSP | 105; N=100 participants | 26 | 11 | 9 | 5.21e-04 | 0.020 | 1/f slope across all seven sources; Relative beta in RAI and LAI | Corrected direct syndrome PSD effect: bvFTD shows more negative 1/f slope and lower beta power than PSP, but individual-level PSD classification is modest. |

| Contrast | No. features | p<0.05 | p<0.01 | q<0.05 | min p | min q | Dominant families | Interpretation |
|---|---|---|---|---|---|---|---|---|
| CON vs pooled patients | 105 | 5 | 0 | 0 | 0.029 | 0.446 | Beta residual max (7 top-ranked); Alpha residual max (2); Relative theta (1) | Weak common disease-control axis; no corrected features after pooling 45 bvFTD and 55 PSP, and pooled PSD classification is weak (best AUC 0.591). |

#### B.4. Contrast-specific statistical results

##### B.4.1. Controls versus bvFTD

This was the only PSD feature contrast with effects surviving Benjamini-Hochberg correction at q_BH < 0.05. The corrected pattern was concentrated in increased alpha-phase/beta-amplitude PAC in bvFTD and reduced beta residual structure, relative beta power, and spectral entropy in bvFTD.

**Table B4. CON vs bvFTD: FDR-significant PSD features (q_BH < 0.05). A = CON; B = bvFTD.**

| Rank | Feature family | Source | Higher mean | Mean A | Mean B | p_Welch | q_BH | Cohen's d |
|---|---|---|---|---|---|---|---|---|
| 1 | PAC alpha-phase/beta-amp | RAI | bvFTD | 1.87e-03 | 2.35e-03 | 7.39e-04 | 0.029 | -0.66 |
| 2 | PAC alpha-phase/beta-amp | LAI | bvFTD | 1.89e-03 | 2.37e-03 | 0.001 | 0.029 | -0.65 |
| 3 | PAC alpha-phase/beta-amp | RAG | bvFTD | 1.96e-03 | 2.45e-03 | 0.002 | 0.029 | -0.61 |
| 4 | Beta residual max | RAI | CON | 1.320 | 0.898 | 9.36e-04 | 0.029 | 0.60 |
| 5 | PAC alpha-phase/beta-amp | PCC | bvFTD | 1.98e-03 | 2.45e-03 | 0.003 | 0.029 | -0.59 |
| 6 | PAC alpha-phase/beta-amp | ACC | bvFTD | 1.85e-03 | 2.30e-03 | 0.003 | 0.029 | -0.59 |
| 7 | PAC alpha-phase/beta-amp | MPFC | bvFTD | 1.83e-03 | 2.26e-03 | 0.004 | 0.029 | -0.57 |
| 8 | Beta residual max | RAG | CON | 1.347 | 0.937 | 0.002 | 0.029 | 0.57 |
| 9 | Beta residual max | LAI | CON | 1.301 | 0.898 | 0.002 | 0.029 | 0.57 |
| 10 | Beta residual max | ACC | CON | 1.357 | 0.925 | 0.002 | 0.029 | 0.56 |
| 11 | Beta residual max | PCC | CON | 1.324 | 0.939 | 0.003 | 0.029 | 0.53 |
| 12 | Beta residual max | LAG | CON | 1.351 | 0.951 | 0.004 | 0.029 | 0.53 |
| 13 | Relative beta | RAI | CON | 0.057 | 0.022 | 0.001 | 0.029 | 0.53 |
| 14 | Relative beta | LAI | CON | 0.059 | 0.022 | 0.003 | 0.029 | 0.47 |
| 15 | PAC alpha-phase/beta-amp | LAG | bvFTD | 1.95e-03 | 2.41e-03 | 0.005 | 0.032 | -0.57 |
| 16 | Beta residual max | MPFC | CON | 1.359 | 0.950 | 0.005 | 0.032 | 0.52 |
| 17 | Spectral entropy | ACC | CON | 0.494 | 0.474 | 0.006 | 0.034 | 0.52 |
| 18 | Relative beta | RAG | CON | 0.055 | 0.023 | 0.006 | 0.037 | 0.44 |
| 19 | Spectral entropy | LAI | CON | 0.482 | 0.461 | 0.007 | 0.041 | 0.52 |
| 20 | Spectral entropy | MPFC | CON | 0.494 | 0.474 | 0.009 | 0.050 | 0.50 |

The sign of Cohen's d follows Group A minus Group B. Negative d values for PAC therefore indicate higher PAC in bvFTD, whereas positive d values for beta residual, relative beta, and spectral entropy indicate higher values in controls and lower values in bvFTD.

##### B.4.2. Controls versus PSP

The CON-PSP feature table used 74 controls and 55 PSP participants. No PSD feature survived FDR correction. One feature, ACC 1/f slope, reached uncorrected p < 0.05, but the adjusted q-value was 0.763, so it is treated as a weak trend rather than a reliable single-feature marker. The top-ranked features were dominated by 1/f slope across ACC, MPFC, RAG, LAG and LAI, followed by beta peak frequency across posterior and salience/default-mode sources. This supports a subtle distributed aperiodic/beta-peak trend rather than a corrected PSP-control PSD signature. The CON-PSP PSD classifier was weak: the best operating point was AdaBoost with the D5 panel (accuracy 0.589; sensitivity 0.509; specificity 0.649; precision 0.519; F1-score 0.514; MCC 0.158; AUC 0.582).

**Table B5. CON vs PSP: top ten ranked PSD features. No feature survived FDR correction.**

| Rank | Feature family | Source | Higher mean | Mean A | Mean B | p_Welch | q_BH | Cohen's d |
|---|---|---|---|---|---|---|---|---|
| 1 | 1/f slope | ACC | PSP | -3.313 | -3.056 | 0.041 | 0.763 | -0.374 |
| 2 | 1/f slope | MPFC | PSP | -3.284 | -3.049 | 0.059 | 0.763 | -0.341 |
| 3 | 1/f slope | RAG | PSP | -3.289 | -3.054 | 0.063 | 0.763 | -0.340 |
| 4 | 1/f slope | LAG | PSP | -3.291 | -3.056 | 0.063 | 0.763 | -0.340 |
| 5 | 1/f slope | LAI | PSP | -3.275 | -3.052 | 0.073 | 0.763 | -0.328 |
| 6 | Beta peak frequency | LAG | PSP | 18.086 | 18.736 | 0.088 | 0.763 | -0.321 |
| 7 | Beta peak frequency | RAI | PSP | 18.055 | 18.686 | 0.089 | 0.763 | -0.320 |

| Rank | Feature family | Source | Higher mean | Mean A | Mean B | p_Welch | q_BH | Cohen's d |
|---|---|---|---|---|---|---|---|---|
| 8 | Beta peak frequency | MPFC | PSP | 18.103 | 18.751 | 0.091 | 0.763 | -0.317 |
| 9 | Beta peak frequency | ACC | PSP | 18.101 | 18.753 | 0.091 | 0.763 | -0.318 |
| 10 | Beta peak frequency | LAI | PSP | 18.105 | 18.734 | 0.096 | 0.763 | -0.314 |

##### B.4.3. bvFTD versus PSP

The direct syndrome PSD difference is dominated by aperiodic spectral organisation: 1/f slope survived FDR correction across all seven SN-DMN sources. Numerically, PSP had the higher, less negative 1/f slope, whereas bvFTD showed the more negative slope. Relative beta power in RAI and LAI also survived FDR correction, with PSP showing higher beta power than bvFTD; relative beta in RAG was the next-ranked trend. In general bvFTD shows a more globally redistributed resting spectral state, with stronger aperiodic involvement and weaker beta-band organisation than PSP.

The individual-level PSD classification is modest despite the clearer univariate aperiodic/beta pattern. The highest AUC was obtained by a linear SVM using ten-feature panels dominated by 1/f slope and relative beta (AUC 0.652; accuracy 0.610; sensitivity 0.533; specificity 0.673; F1-score 0.552; MCC 0.208). The strongest operating point by accuracy and MCC was AdaBoost using the D5 panel (accuracy 0.640; sensitivity 0.533; specificity 0.727; precision 0.615; F1-score 0.571; MCC 0.266; AUC 0.621). Thus, the univariate group-level PSD syndrome effect is physiologically clearer than its individual-level classification coherence.

**Table B6. bvFTD-PSP: top ten ranked PSD features from the univariate analysis. Positive Cohen's d values indicate higher values in PSP than bvFTD.**

| Rank | Feature family | Source | Higher mean | Mean A | Mean B | p_Welch | q_BH | Cohen's d |
|---|---|---|---|---|---|---|---|---|
| 1 | 1/f slope | ACC | PSP | -3.056 | -3.542 | 5.21e-04 | 0.020 | 0.710 |
| 2 | 1/f slope | LAG | PSP | -3.056 | -3.538 | 7.50e-04 | 0.020 | 0.690 |
| 3 | 1/f slope | LAI | PSP | -3.052 | -3.511 | 8.29e-04 | 0.020 | 0.682 |
| 4 | 1/f slope | MPFC | PSP | -3.049 | -3.510 | 8.80e-04 | 0.020 | 0.683 |
| 5 | 1/f slope | RAG | PSP | -3.054 | -3.526 | 9.59e-04 | 0.020 | 0.677 |
| 6 | 1/f slope | RAI | PSP | -3.121 | -3.557 | 0.001 | 0.023 | 0.649 |
| 7 | 1/f slope | PCC | PSP | -3.061 | -3.515 | 0.002 | 0.023 | 0.645 |
| 8 | Relative beta | RAI | PSP | 0.059 | 0.022 | 0.003 | 0.034 | 0.583 |
| 9 | Relative beta | LAI | PSP | 0.072 | 0.022 | 0.003 | 0.039 | 0.563 |
| 10 | Relative beta | RAG | PSP | 0.065 | 0.023 | 0.007 | 0.072 | 0.513 |

##### B.4.4. Controls versus pooled patients

The pooled patient-control comparison used 74 controls and 100 pooled patients. It produced five uncorrected effects but no FDR-surviving PSD features. The leading features were beta residual maxima across most sources, with controls showing higher beta residual structure than pooled patients, and a relative-theta trend in RAI with higher values in pooled patients. The pooled contrast therefore supports a weak common disease axis but also confirms that pooling bvFTD and PSP dilutes syndrome-specific PSD structure.

The pooled-patient PSD classification result is also weak. The most defensible operating point was obtained with an RBF-SVM using a compact five-feature beta-residual/theta panel: beta residual maximum in RAI, RAG, MPFC and ACC, plus relative theta power in RAI. This model achieved accuracy 0.586, sensitivity 0.660, specificity 0.486, precision 0.635, F1-score 0.647, MCC 0.148 and AUC 0.591 (TP 66, TN 36, FP 38, FN 34; N = 174). A sensitivity-biased AdaBoost solution using the same compact panel reached higher apparent accuracy (0.621) and sensitivity (0.880), but low specificity (0.270) and poor AUC (0.476), so it is not treated as stronger physiological evidence. The univariate and classification results converge on the same conclusion: the pooled PSD axis is shallow and should not be interpreted as a robust individual-level marker.

**Table B7. CON-pooled patients: top ten ranked PSD features. No feature survived FDR correction.**

| Rank | Feature family | Source | Higher mean | Mean A | Mean B | p_Welch | q_BH | Cohen's d |
|---|---|---|---|---|---|---|---|---|
| 1 | Beta residual max | RAI | CON | 1.320 | 1.070 | 0.029 | 0.446 | 0.344 |

| Rank | Feature family | Source | Higher mean | Mean A | Mean B | p_Welch | q_BH | Cohen's d |
|---|---|---|---|---|---|---|---|---|
| 2 | Relative theta | RAI | Patient | 0.161 | 0.201 | 0.031 | 0.446 | -0.335 |
| 3 | Beta residual max | RAG | CON | 1.347 | 1.105 | 0.038 | 0.446 | 0.325 |
| 4 | Beta residual max | MPFC | CON | 1.359 | 1.109 | 0.044 | 0.446 | 0.316 |
| 5 | Beta residual max | ACC | CON | 1.357 | 1.107 | 0.046 | 0.446 | 0.315 |
| 6 | Beta residual max | LAG | CON | 1.351 | 1.121 | 0.057 | 0.446 | 0.296 |
| 7 | Alpha residual max | LAG | CON | 1.998 | 1.804 | 0.058 | 0.446 | 0.294 |
| 8 | Alpha residual max | PCC | CON | 2.028 | 1.824 | 0.060 | 0.446 | 0.292 |
| 9 | Beta residual max | PCC | CON | 1.324 | 1.107 | 0.066 | 0.446 | 0.286 |
| 10 | Beta residual max | LAI | CON | 1.301 | 1.085 | 0.066 | 0.446 | 0.286 |

Appendix B: PSD statistical test results

#### B.5. Interpretation

**Table B8. statistical test interpretation.**

| Contrast | Statistical status | Manuscript implication |
|---|---|---|
| CON vs bvFTD | Corrected group-level PSD effect. | bvFTD shows the clearest patient-control spectral reorganisation: higher PAC and lower beta residual/relative beta/spectral entropy. This justifies giving this contrast the strongest patient-control PSD interpretation in the Results. |
| CON vs PSP | No corrected univariate PSD effect; weak classifier. | The PSP results support a subtle PSP-control spectral phenotype. One 1/f-slope feature reached uncorrected $p<0.05$, but no feature survived FDR correction and the best PSD classifier reached only accuracy 0.589, MCC 0.158 and AUC 0.582. |
| bvFTD vs PSP | Corrected direct syndrome PSD effect, but modest individual-level PSD coherence. | The PSP univariate statistical analysis shows FDR-corrected differences in widespread 1/f slope and relative beta power, supporting an aperiodic/beta syndrome axis. However, classification is modest (highest AUC 0.652; best accuracy 0.640), so the result should be framed as physiological structure rather than a robust standalone individual marker. |
| CON vs pooled patients | Weak uncorrected common disease axis; weak individual-level PSD classification. | Pooling identifies reduced beta residual structure and RAI theta as shared trends, but the absence of corrected effects and the weak classifier (accuracy 0.586, MCC 0.148, AUC 0.591) support the conclusion that pooling dilutes syndrome-specific physiology. |

The univariate PSD tests show robust evidence for bvFTD-control spectral reorganization and direct bvFTD-versus-PSP effects centred on widespread 1/f slope and relative beta power. In contrast, CON-PSP remains weak at the univariate level, with only one uncorrected trend and no FDR-surviving features, and the PSD classifier confirms weak individual-level coherence. The pooled patient-control comparison also is weak where no pooled PSD feature survived FDR correction and the best threshold-independent pooled PSD classifier reached only accuracy 0.586, MCC 0.148 and AUC 0.591.

### Appendix C. PSD- and DCM-based classification results

The reported result in the paper is given in Table 1C.

Table 1C. *Extracted features, physiological meaning, and group-to-individual interpretation. They are intended to aid physiological interpretation of group effects and group-to-individual translation, not to define diagnostic biomarkers.*

| Contrast | Domain | Best reported model / panel | Performance | Interpretation |
|---|---|---|---|---|
| CON vs bvFTD | Clinical | Multivariate clinical model; ACE-R and FAB strong individually | Multivariate accuracy 0.915; ACE-R accuracy 0.913; FAB accuracy 0.869 | Clinical impairment is strongly and individually expressed. |
| CON vs bvFTD | PSD | Random forest, S5/P5 panels dominated by PAC and beta-residual features in right anterior insula (RAI), left anterior insula (LAI), and ACC | Best accuracy 0.639; F1 0.494; MCC 0.215; AUC up to 0.627 | Clear group spectral phenotype but modest individual-level coherence. |
| CON vs bvFTD | DCM | Linear SVM, ten-parameter panel | Accuracy 0.748; specificity 0.878; MCC 0.446; AUC 0.756 | Distributed timing, local-gain and AMPA-coupling parameters show moderate subject-level coherence. |
| CON vs PSP | Clinical | Multivariate clinical model; ACE-R and FAB strong individually | Multivariate accuracy 0.936; ACE-R accuracy 0.904; FAB accuracy 0.863; MMSE accuracy 0.775 | PSP-control separation is clinically strong and individually coherent across global cognitive and frontal-executive scores. |
| CON vs PSP | PSD | AdaBoost, D5 panel dominated by 1/f slope in ACC, MPFC, LAG, and RAG plus relative gamma in LAG | Accuracy 0.589; sensitivity 0.509; specificity 0.649; F1 0.514; MCC 0.158; AUC 0.582 | PSD classifier performance is weak; focal spectral effects do not translate into moderate individual-level coherence. |
| CON vs PSP | DCM | Linear SVM, P10 DCM panel | Accuracy 0.698; sensitivity 0.491; specificity 0.851; precision 0.711; F1 0.581; MCC 0.371; AUC 0.770 | Moderate multivariate DCM coherence; high control specificity but lower PSP sensitivity; no single DCM parameter survived FDR. |
| CON vs pooled patients | Clinical | ACE-R and FAB strong; multivariate clinical model lower than single ACE-R | ACE-R 0.907; FAB 0.855; multivariate 0.835 | Shared clinical disease burden is individually coherent. |
| CON vs pooled patients | PSD | RBF-SVM, compact five-feature beta-residual/theta panel | Accuracy 0.586; sensitivity 0.660; specificity 0.486; F1 0.647; MCC 0.148; AUC 0.591 | Pooled PSD coherence is weak; no derived PSD feature survived FDR correction. |
| CON vs pooled patients | DCM | Random Forest P10/stability ten-parameter panel by accuracy/F1/MCC; linear SVM five-parameter panel by AUC | Accuracy up to 0.632; F1 0.692; MCC 0.238; AUC up to 0.648 | A weak-to-modest common gain/coupling disease axis is present but diluted by syndrome heterogeneity. |
| bvFTD vs PSP | Clinical | Multivariate clinical model; PSP strongest single score, with CBI-R, ACE-R, and MMSE also informative | Multivariate accuracy 0.828; PSPRS accuracy 0.843; CBI-R accuracy 0.702; MMSE accuracy 0.691; ACE-R accuracy 0.688 | Direct clinical separation is strong, dominated by PSP motor-functional burden with additional behavioural and cognitive contributions. |
| bvFTD vs PSP | PSD | Linear SVM ten-feature aperiodic/beta panel by AUC; AdaBoost D5 by accuracy/MCC | AUC up to 0.652; best accuracy 0.640; specificity 0.727; F1 0.571; MCC 0.266 | Derived features support an aperiodic/beta syndrome axis, but frequency-bin effects are focal low-frequency and individual-level coherence is modest. |
| bvFTD vs PSP | DCM | Linear SVM, compact five-parameter DCM panel | Accuracy 0.740; PSP sensitivity 0.800; bvFTD specificity 0.667; precision 0.746; F1 0.772; MCC 0.472; AUC 0.798 | Moderate syndrome-level coherence; compact timing/coupling panel involving the RAG AMPA-mediated excitatory time constant, forward NMDA coupling from ACC to MPFC, forward AMPA coupling from LAG to RAG, backward NMDA coupling from MPFC to LAI, and extrinsic conduction/transmission delay. |

The rest of this appendix consolidates detail classification results for each contrast. Metrics are reported from leave-one-out classification. Sensitivity, precision, F1-score, TP, and FN refer to the positive class specified in each contrast table; specificity, TN, and FP refer to the negative class. Note that the feature panels are group-derived summaries of feature importance. The classification values therefore quantify individual-level coherence conditional on those group-derived panels. They are not presented as independently validated prospective diagnostic performance.

##### *Common notation and feature-panel definitions*

PSD source mapping. Ch1 = left anterior insula (LAI), Ch2 = ACC, Ch3 = right anterior insula (RAI), Ch4 = MPFC, Ch5 = PCC, Ch6 = LAG, and Ch7 = RAG.

DCM region indices. LAI = 1, ACC = 2, RAI = 3, MPFC = 4, PCC = 5, LAG = 6, and RAG = 7. Population indices are spiny excitatory = 1, superficial pyramidal = 2, inhibitory = 3, and deep pyramidal = 4.

Feature panels. P5/P10 select the smallest uncorrected p-values; D5/D10 select the largest absolute Cohen d values; C5/C10 use a combined evidence-effect score; S5/S10 use the combined score plus a recurrence/stability bonus.

#### 1. Controls versus bvFTD

##### *PSD-feature classification*

The strongest PSD operating point was Random Forest with the S5 panel (accuracy 0.639; sensitivity 0.467; specificity 0.743; F1-score 0.494; MCC 0.215; AUC 0.617). The selected PSD features were dominated by alpha-phase/beta-amplitude PAC and beta-residual features in salience and default-mode sources, indicating a physiologically coherent but modest individual-level bvFTD spectral phenotype.

**Table 1.1. PSD classification results for controls-bvFTD.**

| Summary criterion | Panel / model | Performance metrics | Confusion matrix |
|---|---|---|---|
| Best balanced result | S5 / RF | Acc 0.639; Sens 0.467; Spec 0.743; Prec 0.525; F1 0.494; MCC 0.215; AUC 0.617 | TP 21; TN 55; FP 19; FN 24 |
| Best accuracy | S5 / RF | Acc 0.639; Sens 0.467; Spec 0.743; Prec 0.525; F1 0.494; MCC 0.215; AUC 0.617 | TP 21; TN 55; FP 19; FN 24 |
| Best available AUC | S5 / RF | Acc 0.639; Sens 0.467; Spec 0.743; Prec 0.525; F1 0.494; MCC 0.215; AUC 0.617 | TP 21; TN 55; FP 19; FN 24 |

**Table 1.2. PSD feature panels used for controls-bvFTD classification.**

| Panel | Rule | Order | Index | Feature family | Source | Channel | Raw workbook feature |
|---|---|---|---|---|---|---|---|
| P10 | 10 smallest uncorrected p-values | 1 | 45 | PAC alpha-phase/beta-amplitude | RAI | Ch3 | PAC_MI_alphaPhase_betaAmp__Ch3 |
| | | 2 | 42 | Beta residual max | RAI | Ch3 | Beta_residual_max_Ch3 |
| | | 3 | 15 | PAC alpha-phase/beta-amplitude | LAI | Ch1 | PAC_MI_alphaPhase_betaAmp__Ch1 |
| | | 4 | 34 | Relative beta | RAI | Ch3 | RelBeta__Ch3 |
| | | 5 | 12 | Beta residual max | LAI | Ch1 | Beta_residual_max__Ch1 |
| | | 6 | 102 | Beta residual max | RAG | Ch7 | Beta_residual_max__Ch7 |
| | | 7 | 27 | Beta residual max | ACC | Ch2 | Beta_residual_max__Ch2 |
| | | 8 | 105 | PAC alpha-phase/beta-amplitude | RAG | Ch7 | PAC_MI_alphaPhase_betaAmp__Ch7 |
| | | 9 | 30 | PAC alpha-phase/beta-amplitude | ACC | Ch2 | PAC_MI_alphaPhase_betaAmp__Ch2 |
| | | 10 | 75 | PAC alpha-phase/beta-amplitude | PCC | Ch5 | PAC_MI_alphaPhase_betaAmp__Ch5 |
| D5 | 5 largest absolute | 1 | 45 | PAC alpha- | RAI | Ch3 | PAC_MI_alphaPhase_betaAmp__Ch3 |

| Panel | Rule | Order | Index | Feature family | Source | Channel | Raw workbook feature |
|---|---|---|---|---|---|---|---|
| | Cohen's d values | | | phase/beta-amplitude | | | |
| | | 2 | 15 | PAC alpha-phase/beta-amplitude | LAI | Ch1 | PAC_MI_alphaPhase_betaAmp__Ch1 |
| | | 3 | 105 | PAC alpha-phase/beta-amplitude | RAG | Ch7 | PAC_MI_alphaPhase_betaAmp__Ch7 |
| | | 4 | 42 | Beta residual max | RAI | Ch3 | Beta_residual_max_Ch3 |
| | | 5 | 75 | PAC alpha-phase/beta-amplitude | PCC | Ch5 | PAC_MI_alphaPhase_betaAmp__Ch5 |
| D10 | 10 largest absolute Cohen's d values | 1 | 45 | PAC alpha-phase/beta-amplitude | RAI | Ch3 | PAC_MI_alphaPhase_betaAmp__Ch3 |
| | | 2 | 15 | PAC alpha-phase/beta-amplitude | LAI | Ch1 | PAC_MI_alphaPhase_betaAmp__Ch1 |
| | | 3 | 105 | PAC alpha-phase/beta-amplitude | RAG | Ch7 | PAC_MI_alphaPhase_betaAmp__Ch7 |
| | | 4 | 42 | Beta residual max | RAI | Ch3 | Beta_residual_max__Ch3 |
| | | 5 | 75 | PAC alpha-phase/beta-amplitude | PCC | Ch5 | PAC_MI_alphaPhase_betaAmp__Ch5 |
| | | 6 | 30 | PAC alpha-phase/beta-amplitude | ACC | Ch2 | PAC_MI_alphaPhase_betaAmp__Ch2 |
| | | 7 | 60 | PAC alpha-phase/beta-amplitude | MPFC | Ch4 | PAC_MI_alphaPhase_betaAmp__Ch4 |
| | | 8 | 102 | Beta residual max | RAG | Ch7 | Beta_residual_max__Ch7 |
| | | 9 | 12 | Beta residual max | LAI | Ch1 | Beta_residual_max__Ch1 |
| | | 10 | 90 | PAC alpha-phase/beta-amplitude | LAG | Ch6 | PAC_MI_alphaPhase_betaAmp__Ch6 |
| C5 | Top 5 combined evidence-effect score | 1 | 45 | PAC alpha-phase/beta-amplitude | RAI | Ch3 | PAC_MI_alphaPhase_betaAmp__Ch3 |
| | | 2 | 15 | PAC alpha-phase/beta-amplitude | LAI | Ch1 | PAC_MI_alphaPhase_betaAmp__Ch1 |
| | | 3 | 42 | Beta residual max | RAI | Ch3 | Beta_residual_max_Ch3 |
| | | 4 | 105 | PAC alpha-phase/beta-amplitude | RAG | Ch7 | PAC_MI_alphaPhase_betaAmp__Ch7 |
| | | 5 | 12 | Beta residual max | LAI | Ch1 | Beta_residual_max__Ch1 |
| C10 | Top 10 combined evidence-effect score | 1 | 45 | PAC alpha-phase/beta-amplitude | RAI | Ch3 | PAC_MI_alphaPhase_betaAmp__Ch3 |
| | | 2 | 15 | PAC alpha-phase/beta-amplitude | LAI | Ch1 | PAC_MI_alphaPhase_betaAmp__Ch1 |
| | | 3 | 42 | Beta residual max | RAI | Ch3 | Beta_residual_max__Ch3 |
| | | 4 | 105 | PAC alpha-phase/beta-amplitude | RAG | Ch7 | PAC_MI_alphaPhase_betaAmp__Ch7 |
| | | 5 | 12 | Beta residual max | LAI | Ch1 | Beta_residual_max__Ch1 |
| | | 6 | 102 | Beta residual max | RAG | Ch7 | Beta_residual_max__Ch7 |
| | | 7 | 34 | Relative beta | RAI | Ch3 | RelBeta__Ch3 |
| | | 8 | 30 | PAC alpha-phase/beta-amplitude | ACC | Ch2 | PAC_MI_alphaPhase_betaAmp__Ch2 |
| | | 9 | 27 | Beta residual max | ACC | Ch2 | Beta_residual_max__Ch2 |
| | | 10 | 75 | PAC alpha-phase/beta-amplitude | PCC | Ch5 | PAC_MI_alphaPhase_betaAmp__Ch5 |
| S5 | Top 5 combined score plus recurrence bonus | 1 | 45 | PAC alpha-phase/beta-amplitude | RAI | Ch3 | PAC_MI_alphaPhase_betaAmp__Ch3 |
| | | 2 | 15 | PAC alpha-phase/beta-amplitude | LAI | Ch1 | PAC_MI_alphaPhase_betaAmp__Ch1 |
| | | 3 | 42 | Beta residual max | RAI | Ch3 | Beta_residual_max__Ch3 |

| Panel | Rule | Order | Index | Feature family | Source | Channel | Raw workbook feature |
|---|---|---|---|---|---|---|---|
| | | 4 | 12 | Beta residual max | LAI | Ch1 | Beta_residual_max__Ch1 |
| | | 5 | 27 | Beta residual max | ACC | Ch2 | Beta_residual_max__Ch2 |
| S10 | Top 10 combined score plus recurrence bonus | 1 | 45 | PAC alpha-phase/beta-amplitude | RAI | Ch3 | PAC_MI_alphaPhase_betaAmp__Ch3 |
| | | 2 | 15 | PAC alpha-phase/beta-amplitude | LAI | Ch1 | PAC_MI_alphaPhase_betaAmp__Ch1 |
| | | 3 | 42 | Beta residual max | RAI | Ch3 | Beta_residual_max_Ch3 |
| | | 4 | 12 | Beta residual max | LAI | Ch1 | Beta_residual_max__Ch1 |
| | | 5 | 27 | Beta residual max | ACC | Ch2 | Beta_residual_max__Ch2 |
| | | 6 | 102 | Beta residual max | RAG | Ch7 | Beta_residual_max__Ch7 |
| | | 7 | 34 | Relative beta | RAI | Ch3 | RelBeta__Ch3 |
| | | 8 | 105 | PAC alpha-phase/beta-amplitude | RAG | Ch7 | PAC_MI_alphaPhase_betaAmp__Ch7 |
| | | 9 | 30 | PAC alpha-phase/beta-amplitude | ACC | Ch2 | PAC_MI_alphaPhase_betaAmp__Ch2 |
| | | 10 | 75 | PAC alpha-phase/beta-amplitude | PCC | Ch5 | PAC_MI_alphaPhase_betaAmp__Ch5 |

**Table 1.3. Best PSD classifier within each available feature panel.**

| Panel | Best model | Acc. | Sens. | Spec. | Prec. | F1 | MCC | | AUC | TP | TN | FP | FN |
|---|---|---|---|---|---|---|---|---|---|---|---|---|---|
| P10 | AdaBoost | 0.580 | 0.356 | 0.716 | 0.432 | 0.390 | 0.075 | | 0.579 | 16 | 53 | 21 | 29 |
| D5 | AdaBoost | 0.622 | 0.400 | 0.757 | 0.500 | 0.444 | 0.165 | | 0.567 | 18 | 56 | 18 | 27 |
| D10 | Lin-SVM | 0.588 | 0.267 | 0.784 | 0.429 | 0.329 | 0.058 | | 0.559 | 12 | 58 | 16 | 33 |
| C5 | RF | 0.605 | 0.400 | 0.730 | 0.474 | 0.434 | 0.135 | | 0.613 | 18 | 54 | 20 | 27 |
| C10 | AdaBoost | 0.580 | 0.356 | 0.716 | 0.432 | 0.390 | 0.075 | | 0.591 | 16 | 53 | 21 | 29 |
| S5 | RF | 0.639 | 0.467 | 0.743 | 0.525 | 0.494 | 0.215 | | 0.617 | 21 | 55 | 19 | 24 |
| S10 | AdaBoost | 0.580 | 0.356 | 0.716 | 0.432 | 0.390 | 0.075 | | 0.591 | 16 | 53 | 21 | 29 |

**Table 1.4. PSD classifier metrics for controls-bvFTD.**

| Panel | Classifier | Acc. | Sens. | Spec. | Prec. | F1 | MCC | AUC | TP | TN | FP | FN |
|---|---|---|---|---|---|---|---|---|---|---|---|---|
| P10 | Lin-SVM | 0.555 | 0.200 | 0.770 | 0.346 | 0.254 | -0.035 | 0.534 | 9 | 57 | 17 | 36 |
| P10 | RBF-SVM | 0.580 | 0.356 | 0.716 | 0.432 | 0.390 | 0.075 | 0.521 | 16 | 53 | 21 | 29 |
| P10 | RF | 0.563 | 0.333 | 0.703 | 0.405 | 0.366 | 0.038 | 0.593 | 15 | 52 | 22 | 30 |
| P10 | AdaBoost | 0.580 | 0.356 | 0.716 | 0.432 | 0.390 | 0.075 | 0.579 | 16 | 53 | 21 | 29 |
| D5 | Lin-SVM | 0.546 | 0.222 | 0.743 | 0.345 | 0.270 | -0.039 | 0.538 | 10 | 55 | 19 | 35 |
| D5 | RBF-SVM | 0.521 | 0.556 | 0.500 | 0.403 | 0.467 | 0.054 | 0.495 | 25 | 37 | 37 | 20 |
| D5 | RF | 0.555 | 0.333 | 0.689 | 0.395 | 0.361 | 0.023 | 0.570 | 15 | 51 | 23 | 30 |
| D5 | AdaBoost | 0.622 | 0.400 | 0.757 | 0.500 | 0.444 | 0.165 | 0.567 | 18 | 56 | 18 | 27 |
| D10 | Lin-SVM | 0.588 | 0.267 | 0.784 | 0.429 | 0.329 | 0.058 | 0.559 | 12 | 58 | 16 | 33 |
| D10 | RBF-SVM | 0.521 | 0.378 | 0.608 | 0.370 | 0.374 | -0.014 | 0.505 | 17 | 45 | 29 | 28 |
| D10 | RF | 0.546 | 0.267 | 0.716 | 0.364 | 0.308 | -0.019 | 0.590 | 12 | 53 | 21 | 33 |
| D10 | AdaBoost | 0.546 | 0.311 | 0.689 | 0.378 | 0.341 | 0.000 | 0.598 | 14 | 51 | 23 | 31 |
| C5 | Lin-SVM | 0.588 | 0.244 | 0.797 | 0.423 | 0.310 | 0.049 | 0.558 | 11 | 59 | 15 | 34 |
| C5 | RBF-SVM | 0.571 | 0.378 | 0.689 | 0.425 | 0.400 | 0.069 | 0.533 | 17 | 51 | 23 | 28 |
| C5 | RF | 0.605 | 0.400 | 0.730 | 0.474 | 0.434 | 0.135 | 0.613 | 18 | 54 | 20 | 27 |
| C5 | AdaBoost | 0.597 | 0.400 | 0.716 | 0.462 | 0.429 | 0.120 | 0.569 | 18 | 53 | 21 | 27 |
| C10 | Lin-SVM | 0.555 | 0.200 | 0.770 | 0.346 | 0.254 | -0.035 | 0.535 | 9 | 57 | 17 | 36 |
| C10 | RBF-SVM | 0.580 | 0.356 | 0.716 | 0.432 | 0.390 | 0.075 | 0.521 | 16 | 53 | 21 | 29 |
| C10 | RF | 0.563 | 0.333 | 0.703 | 0.405 | 0.366 | 0.038 | 0.594 | 15 | 52 | 22 | 30 |
| C10 | AdaBoost | 0.580 | 0.356 | 0.716 | 0.432 | 0.390 | 0.075 | 0.591 | 16 | 53 | 21 | 29 |
| S5 | Lin-SVM | 0.613 | 0.289 | 0.811 | 0.481 | 0.361 | 0.115 | 0.576 | 13 | 60 | 14 | 32 |
| S5 | RBF-SVM | 0.613 | 0.378 | 0.757 | 0.486 | 0.425 | 0.143 | 0.584 | 17 | 56 | 18 | 28 |
| S5 | RF | 0.639 | 0.467 | 0.743 | 0.525 | 0.494 | 0.215 | 0.617 | 21 | 55 | 19 | 24 |
| S5 | AdaBoost | 0.613 | 0.467 | 0.703 | 0.488 | 0.477 | 0.171 | 0.604 | 21 | 52 | 22 | 24 |
| S10 | Lin-SVM | 0.555 | 0.200 | 0.770 | 0.346 | 0.254 | -0.035 | 0.535 | 9 | 57 | 17 | 36 |
| S10 | RBF-SVM | 0.580 | 0.356 | 0.716 | 0.432 | 0.390 | 0.075 | 0.521 | 16 | 53 | 21 | 29 |
| S10 | RF | 0.555 | 0.333 | 0.689 | 0.395 | 0.361 | 0.023 | 0.594 | 15 | 51 | 23 | 30 |
| S10 | AdaBoost | 0.580 | 0.356 | 0.716 | 0.432 | 0.390 | 0.075 | 0.591 | 16 | 53 | 21 | 29 |

##### *DCM-parameter classification*

The strongest balanced DCM operating point was a linear SVM with a ten-parameter panel (C10/S10/P10-equivalent), with accuracy 0.748, sensitivity 0.533, specificity 0.878, F1-score 0.615, MCC 0.446, and AUC 0.756. The selected parameters combine extrinsic delay, RAG AMPA time constants, superficial-pyramidal capacitance/self-coupling, and AMPA-mediated communication among salience/default-mode regions.

Table 1.5. DCM classification design and label coding for controls versus bvFTD.

| Item | Value |
| --- | --- |
| Contrast | Controls versus bvFTD |
| Sample size | N = 119 total; controls = 74; bvFTD = 45 |
| Label coding | Controls = 0; bvFTD = 1 |
| Positive class for sensitivity / precision / F1 | bvFTD |
| Negative class for specificity | Controls |
| Evaluation scheme | Leave-one-out cross-validation |
| Classifier families | Linear SVM, RBF-SVM, Random Forest, AdaBoost |
| Feature panels | P5/P10, D5/D10, C5/C10, S5/S10 |

**Table 1.6. DCM classification results for controls-bvFTD.**

| Summary criterion | Panel / model | Performance metrics | Confusion matrix |
| --- | --- | --- | --- |
| Balanced result | C10 / Linear SVM | Acc 0.748; Sens 0.533; Spec 0.878; Prec 0.727; F1 0.615; MCC 0.446; AUC 0.756 | TP 24; TN 65; FP 9; FN 21 |
| Highest AUC | D10 / Linear SVM | Acc 0.723; Sens 0.444; Spec 0.892; Prec 0.714; F1 0.548; MCC 0.384; AUC 0.766 | TP 20; TN 66; FP 8; FN 25 |
| Highest accuracy / MCC | C10 / Linear SVM | Acc 0.748; Sens 0.533; Spec 0.878; Prec 0.727; F1 0.615; MCC 0.446; AUC 0.756 | TP 24; TN 65; FP 9; FN 21 |
| Highest F1-score | C10 / Linear SVM | Acc 0.748; Sens 0.533; Spec 0.878; Prec 0.727; F1 0.615; MCC 0.446; AUC 0.756 | TP 24; TN 65; FP 9; FN 21 |

**Table 1.7. Univariate DCM statistical context for controls-bvFTD.**

| Univariate DCM-test context | Value |
| --- | --- |
| Valid DCM parameter tests | 337 |
| Uncorrected p<0.05 | 16 parameters |
| Uncorrected p<0.01 | 4 parameters |
| BH-FDR q<0.05 | 0 parameters |
| Minimum uncorrected p-value | 8.35e-04 |
| Minimum BH-FDR q-value | 0.281 |

**Table 1.8. Selected DCM parameters in each control-bvFTD feature panel.**

| Panel | Selection rule | n | Selected DCM parameters | Dominant parameter families |
| --- | --- | --- | --- | --- |
| P5 | 5 smallest uncorrected p-values | 5 | 1. D(2); 2. T(7,1); 3. CV(2); 4. H(1,1,2); 5. H(2,2,2) | Intrinsic microcircuit coupling (2); Delay (1); Synaptic time constant (1); Membrane capacitance (1) |
| P10 | 10 smallest uncorrected p-values | 10 | 1. D(2); 2. T(7,1); 3. CV(2); 4. H(1,1,2); 5. H(2,2,2); 6. A{1}(3,7); 7. H(2,2,3); 8. A{2}(1,3); 9. A{1}(5,2); 10. H(2,2,4) | Intrinsic microcircuit coupling (4); Extrinsic effective connectivity (3); Delay (1); Synaptic time constant (1); Membrane capacitance (1) |
| D5 | 5 largest absolute Cohen's d values | 5 | 1. D(2); 2. H(1,1,2); 3. H(2,2,2); 4. CV(2); 5. A{1}(3,7) | Intrinsic microcircuit coupling (2); Delay (1); Membrane capacitance (1); Extrinsic effective connectivity (1) |
| D10 | 10 largest absolute Cohen's d values | 10 | 1. D(2); 2. H(1,1,2); 3. H(2,2,2); 4. CV(2); 5. A{1}(3,7); 6. A{1}(5,2); 7. T(7,1); 8. H(2,2,3); 9. AN{1}(5,2); 10. H(2,2,4) | Intrinsic microcircuit coupling (4); Extrinsic effective connectivity (3); Delay (1); Membrane capacitance (1); Synaptic time constant (1) |
| C5 | Top 5 combined evidence-effect score | 5 | 1. D(2); 2. T(7,1); 3. H(1,1,2); 4. CV(2); 5. H(2,2,2) | Intrinsic microcircuit coupling (2); Delay (1); Synaptic time constant (1); Membrane capacitance (1) |
| C10 | Top 10 combined evidence-effect score | 10 | 1. D(2); 2. T(7,1); 3. H(1,1,2); 4. CV(2); 5. H(2,2,2); 6. A{1}(3,7); 7. H(2,2,3); 8. A{1}(5,2); 9. A{2}(1,3); 10. H(2,2,4) | Intrinsic microcircuit coupling (4); Extrinsic effective connectivity (3); Delay (1); Synaptic time constant (1); Membrane capacitance (1) |

| Panel | Selection rule | n | Selected DCM parameters | Dominant parameter families |
|---|---|---|---|---|
| S5 | Top 5 combined score plus recurrence bonus | 5 | 1. D(2); 2. T(7,1); 3. H(1,1,2); 4. CV(2); 5. H(2,2,2) | Intrinsic microcircuit coupling (2); Delay (1); Synaptic time constant (1); Membrane capacitance (1) |
| S10 | Top 10 combined score plus recurrence bonus | 10 | 1. D(2); 2. T(7,1); 3. H(1,1,2); 4. CV(2); 5. H(2,2,2); 6. A{1}(3,7); 7. H(2,2,3); 8. A{1}(5,2); 9. H(2,2,4); 10. A{2}(1,3) | Intrinsic microcircuit coupling (4); Extrinsic effective connectivity (3); Delay (1); Synaptic time constant (1); Membrane capacitance (1) |

##### Table 1.9. Physiological interpretation of DCM parameters selected across panels.

| Parameter | Family | Physiological meaning | Higher mean | p_Welch | q_BH | Cohen d |
|---|---|---|---|---|---|---|
| D(2) | Delay | extrinsic conduction/transmission delay | CON | 0.001 | 0.281 | 0.64 |
| T(7,1) | Synaptic time constant | AMPA-mediated excitatory synaptic time constant in RAG | CON | 0.003 | 0.520 | 0.51 |
| CV(2) | Membrane capacitance | membrane capacitance for superficial pyramidal population | bvFTD | 0.006 | 0.657 | -0.55 |
| H(1,1,2) | Intrinsic microcircuit coupling | intrinsic coupling in ACC: spiny excitatory to spiny excitatory | CON | 0.008 | 0.657 | 0.60 |
| H(2,2,2) | Intrinsic microcircuit coupling | intrinsic coupling in ACC: superficial pyramidal to superficial pyramidal | CON | 0.012 | 0.730 | 0.57 |
| A{1}(3,7) | Extrinsic effective connectivity | forward AMPA coupling from RAG to RAI | bvFTD | 0.015 | 0.730 | -0.54 |
| H(2,2,3) | Intrinsic microcircuit coupling | intrinsic coupling in RAI: superficial pyramidal to superficial pyramidal | CON | 0.015 | 0.730 | 0.51 |
| A{2}(1,3) | Extrinsic effective connectivity | backward AMPA coupling from RAI to LAI | CON | 0.020 | 0.787 | 0.44 |
| A{1}(5,2) | Extrinsic effective connectivity | forward AMPA coupling from ACC to PCC | bvFTD | 0.021 | 0.787 | -0.51 |
| H(2,2,4) | Intrinsic microcircuit coupling | intrinsic coupling in MPFC: superficial pyramidal to superficial pyramidal | CON | 0.026 | 0.839 | 0.46 |
| AN{1}(5,2) | Extrinsic effective connectivity | forward NMDA coupling from ACC to PCC | | | | |

##### Table 1.10. Best-performing DCM classifier within each feature panel.

| Panel | Best model by AUC | Accuracy | Sensitivity | Specificity | Precision | F1 | MCC | AUC | TP | TN | FP | FN |
|---|---|---|---|---|---|---|---|---|---|---|---|---|
| P5 | Linear SVM | 0.697 | 0.422 | 0.865 | 0.655 | 0.514 | 0.324 | 0.763 | 19 | 64 | 10 | 26 |
| P10 | Linear SVM | 0.748 | 0.533 | 0.878 | 0.727 | 0.615 | 0.446 | 0.756 | 24 | 65 | 9 | 21 |
| D5 | Linear SVM | 0.731 | 0.467 | 0.892 | 0.724 | 0.568 | 0.405 | 0.761 | 21 | 66 | 8 | 24 |
| D10 | Linear SVM | 0.723 | 0.444 | 0.892 | 0.714 | 0.548 | 0.384 | 0.766 | 20 | 66 | 8 | 25 |
| C5 | Linear SVM | 0.697 | 0.422 | 0.865 | 0.655 | 0.514 | 0.324 | 0.763 | 19 | 64 | 10 | 26 |
| C10 | Linear SVM | 0.748 | 0.533 | 0.878 | 0.727 | 0.615 | 0.446 | 0.756 | 24 | 65 | 9 | 21 |
| S5 | Linear SVM | 0.697 | 0.422 | 0.865 | 0.655 | 0.514 | 0.324 | 0.763 | 19 | 64 | 10 | 26 |
| S10 | Linear SVM | 0.748 | 0.533 | 0.878 | 0.727 | 0.615 | 0.446 | 0.756 | 24 | 65 | 9 | 21 |

##### Table 1.11. Full DCM classifier metrics for controls versus bvFTD.

| Panel | Model | Accuracy | Sensitivity | Specificity | Precision | F1 | MCC | AUC | TP | TN | FP | FN |
|---|---|---|---|---|---|---|---|---|---|---|---|---|
| P5 | Linear SVM | 0.697 | 0.422 | 0.865 | 0.655 | 0.514 | 0.324 | 0.763 | 19 | 64 | 10 | 26 |
| P5 | RBF-SVM | 0.571 | 0.689 | 0.500 | 0.456 | 0.549 | 0.185 | 0.541 | 31 | 37 | 37 | 14 |
| P5 | Random Forest | 0.664 | 0.422 | 0.811 | 0.576 | 0.487 | 0.252 | 0.679 | 19 | 60 | 14 | 26 |
| P5 | AdaBoost | 0.647 | 0.422 | 0.784 | 0.543 | 0.475 | 0.219 | 0.649 | 19 | 58 | 16 | 26 |
| P10 | Linear SVM | 0.748 | 0.533 | 0.878 | 0.727 | 0.615 | 0.446 | 0.756 | 24 | 65 | 9 | 21 |
| P10 | RBF-SVM | 0.513 | 0.689 | 0.405 | 0.413 | 0.517 | 0.095 | 0.618 | 31 | 30 | 44 | 14 |
| P10 | Random Forest | 0.723 | 0.511 | 0.851 | 0.676 | 0.582 | 0.389 | 0.685 | 23 | 63 | 11 | 22 |
| P10 | AdaBoost | 0.723 | 0.511 | 0.851 | 0.676 | 0.582 | 0.389 | 0.733 | 23 | 63 | 11 | 22 |
| D5 | Linear SVM | 0.731 | 0.467 | 0.892 | 0.724 | 0.568 | 0.405 | 0.761 | 21 | 66 | 8 | 24 |
| D5 | RBF-SVM | 0.580 | 0.689 | 0.514 | 0.463 | 0.554 | 0.198 | 0.596 | 31 | 38 | 36 | 14 |
| D5 | Random Forest | 0.723 | 0.489 | 0.865 | 0.688 | 0.571 | 0.387 | 0.689 | 22 | 64 | 10 | 23 |
| D5 | AdaBoost | 0.639 | 0.444 | 0.757 | 0.526 | 0.482 | 0.209 | 0.648 | 20 | 56 | 18 | 25 |
| D10 | Linear SVM | 0.723 | 0.444 | 0.892 | 0.714 | 0.548 | 0.384 | 0.766 | 20 | 66 | 8 | 25 |
| D10 | RBF-SVM | 0.504 | 0.667 | 0.405 | 0.405 | 0.504 | 0.072 | 0.584 | 30 | 30 | 44 | 15 |
| D10 | Random Forest | 0.681 | 0.467 | 0.811 | 0.600 | 0.525 | 0.295 | 0.711 | 21 | 60 | 14 | 24 |
| D10 | AdaBoost | 0.723 | 0.467 | 0.878 | 0.700 | 0.560 | 0.385 | 0.673 | 21 | 65 | 9 | 24 |
| C5 | Linear SVM | 0.697 | 0.422 | 0.865 | 0.655 | 0.514 | 0.324 | 0.763 | 19 | 64 | 10 | 26 |
| C5 | RBF-SVM | 0.571 | 0.689 | 0.500 | 0.456 | 0.549 | 0.185 | 0.541 | 31 | 37 | 37 | 14 |
| C5 | Random Forest | 0.664 | 0.422 | 0.811 | 0.576 | 0.487 | 0.252 | 0.676 | 19 | 60 | 14 | 26 |
| C5 | AdaBoost | 0.655 | 0.422 | 0.797 | 0.559 | 0.481 | 0.236 | 0.660 | 19 | 59 | 15 | 26 |
| C10 | Linear SVM | 0.748 | 0.533 | 0.878 | 0.727 | 0.615 | 0.446 | 0.756 | 24 | 65 | 9 | 21 |

| Panel | Model | Accuracy | Sensitivity | Specificity | Precision | F1 | MCC | AUC | TP | TN | FP | FN |
|---|---|---|---|---|---|---|---|---|---|---|---|---|
| C10 | RBF-SVM | 0.513 | 0.689 | 0.405 | 0.413 | 0.517 | 0.095 | 0.618 | 31 | 30 | 44 | 14 |
| C10 | Random Forest | 0.723 | 0.511 | 0.851 | 0.676 | 0.582 | 0.389 | 0.682 | 23 | 63 | 11 | 22 |
| C10 | AdaBoost | 0.723 | 0.533 | 0.838 | 0.667 | 0.593 | 0.392 | 0.719 | 24 | 62 | 12 | 21 |
| S5 | Linear SVM | 0.697 | 0.422 | 0.865 | 0.655 | 0.514 | 0.324 | 0.763 | 19 | 64 | 10 | 26 |
| S5 | RBF-SVM | 0.571 | 0.689 | 0.500 | 0.456 | 0.549 | 0.185 | 0.541 | 31 | 37 | 37 | 14 |
| S5 | Random Forest | 0.664 | 0.422 | 0.811 | 0.576 | 0.487 | 0.252 | 0.676 | 19 | 60 | 14 | 26 |
| S5 | AdaBoost | 0.655 | 0.422 | 0.797 | 0.559 | 0.481 | 0.236 | 0.660 | 19 | 59 | 15 | 26 |
| S10 | Linear SVM | 0.748 | 0.533 | 0.878 | 0.727 | 0.615 | 0.446 | 0.756 | 24 | 65 | 9 | 21 |
| S10 | RBF-SVM | 0.513 | 0.689 | 0.405 | 0.413 | 0.517 | 0.095 | 0.618 | 31 | 30 | 44 | 14 |
| S10 | Random Forest | 0.714 | 0.511 | 0.838 | 0.657 | 0.575 | 0.371 | 0.681 | 23 | 62 | 12 | 22 |
| S10 | AdaBoost | 0.723 | 0.533 | 0.838 | 0.667 | 0.593 | 0.392 | 0.733 | 24 | 62 | 12 | 21 |

##### *Contrast-level interpretation*

Controls-bvFTD shows the clearest physiological coherence among the patient-control contrasts. PSD features describe altered cross-frequency organisation and reduced beta residual structure, but the PSD classifier remains modest. DCM provides stronger coherence, with a linear additive pattern across timing, local gain/capacitance, and AMPA-mediated network communication. The contrast supports a distributed bvFTD physiological phenotype with partial but not complete individual-level separation.

#### 2. Controls versus PSP

##### *PSD-feature classification*

The strongest PSD operating point was AdaBoost with the D5 panel, giving accuracy 0.589, sensitivity 0.509, specificity 0.649, precision 0.519, F1-score 0.514, MCC 0.158, and AUC 0.582. The PSD analysis were dominated by 1/f slope across LAI, ACC, MPFC, LAG and RAG, with smaller contributions from beta peak frequency and relative gamma power in LAG. This agrees with the frequency-bin result, where CON-PSP binwise effects were focal low-frequency effects in LAI and RAG and no derived PSD feature survived FDR correction.

**Table 2.1. PSD classification design and label coding for controls -PSP.**

| Item | Value |
|---|---|
| Contrast | Controls versus PSP |
| Sample size | N = 129 total; controls = 74; PSP = 55 |
| Label coding | Controls = 0; PSP = 1 |
| Positive class for sensitivity / precision / F1 | PSP |
| Negative class for specificity | Controls |
| Evaluation scheme | Leave-one-out cross-validation |
| Classifier families | Linear SVM, RBF-SVM, Random Forest, AdaBoost |
| Feature panels | P5/P10, D5/D10, C5/C10, S5/S10 |

**Table 2.2. Best PSD classifier within each feature panel.**

| Panel | Best model by AUC | Accuracy | Sensitivity | Specificity | Precision | F1 | MCC | AUC |
|---|---|---|---|---|---|---|---|---|
| P5 | RBF-SVM | 0.519 | 0.327 | 0.662 | 0.419 | 0.367 | -0.011 | 0.533 |
| P10 | Lin-SVM | 0.581 | 0.255 | 0.824 | 0.519 | 0.341 | 0.096 | 0.524 |
| D5 | AdaBoost | 0.589 | 0.509 | 0.649 | 0.519 | 0.514 | 0.158 | 0.582 |
| D10 | Lin-SVM | 0.574 | 0.200 | 0.851 | 0.500 | 0.286 | 0.068 | 0.538 |
| C5 | RBF-SVM | 0.519 | 0.327 | 0.662 | 0.419 | 0.367 | -0.011 | 0.533 |
| C10 | Lin-SVM | 0.574 | 0.200 | 0.851 | 0.500 | 0.286 | 0.068 | 0.538 |
| S5 | RBF-SVM | 0.519 | 0.327 | 0.662 | 0.419 | 0.367 | -0.011 | 0.533 |
| S10 | Lin-SVM | 0.574 | 0.200 | 0.851 | 0.500 | 0.286 | 0.068 | 0.538 |

**Table 2.3. PSD feature panels used for controls -PSP classification.**

| Panel | Selection rule | n | Selected PSD features and source regions |
|---|---|---|---|
| P5 | 5 smallest uncorrected p-values | 5 | 1. 1/f slope (ACC); 2. 1/f slope (MPFC); 3. 1/f slope (RAG); 4. 1/f slope (LAG); 5. 1/f slope (LAI) |
| P10 | 10 smallest uncorrected p-values | 10 | 1. 1/f slope (ACC); 2. 1/f slope (MPFC); 3. 1/f slope (RAG); 4. 1/f slope (LAG); 5. 1/f slope (LAI); 6. |

| Panel | Selection rule | n | Selected PSD features and source regions |
|---|---|---|---|
| | | | Beta peak frequency (LAG); 7. Beta peak frequency (RAI); 8. Beta peak frequency (MPFC); 9. Beta peak frequency (ACC); 10. Beta peak frequency (LAI) |
| D5 | 5 largest absolute Cohen's d values | 5 | 1. 1/f slope (ACC); 2. Relative gamma power (LAG); 3. 1/f slope (MPFC); 4. 1/f slope (LAG); 5. 1/f slope (RAG) |
| D10 | 10 largest absolute Cohen's d values | 10 | 1. 1/f slope (ACC); 2. Relative gamma power (LAG); 3. 1/f slope (MPFC); 4. 1/f slope (LAG); 5. 1/f slope (RAG); 6. 1/f slope (LAI); 7. Beta peak frequency (LAG); 8. Beta peak frequency (RAI); 9. Beta peak frequency (ACC); 10. Beta peak frequency (MPFC) |
| C5 | Top 5 by combined evidence-effect score | 5 | 1. 1/f slope (ACC); 2. 1/f slope (MPFC); 3. 1/f slope (RAG); 4. 1/f slope (LAG); 5. 1/f slope (LAI) |
| C10 | Top 10 by combined evidence-effect score | 10 | 1. 1/f slope (ACC); 2. 1/f slope (MPFC); 3. 1/f slope (RAG); 4. 1/f slope (LAG); 5. 1/f slope (LAI); 6. Relative gamma power (LAG); 7. Beta peak frequency (LAG); 8. Beta peak frequency (RAI); 9. Beta peak frequency (ACC); 10. Beta peak frequency (MPFC) |
| S5 | Top 5 by combined score plus recurrence bonus | 5 | 1. 1/f slope (ACC); 2. 1/f slope (MPFC); 3. 1/f slope (RAG); 4. 1/f slope (LAG); 5. 1/f slope (LAI) |
| S10 | Top 10 by combined score plus recurrence bonus | 10 | 1. 1/f slope (ACC); 2. 1/f slope (MPFC); 3. 1/f slope (RAG); 4. 1/f slope (LAG); 5. 1/f slope (LAI); 6. Beta peak frequency (RAI); 7. Relative gamma power (LAG); 8. Beta peak frequency (MPFC); 9. Beta peak frequency (LAG); 10. Beta peak frequency (ACC) |

**Table 2.4. Physiological interpretation of selected PSD feature families.**

| Feature family appearing in panels | Regions selected | Panel appearances | Interpretation for this contrast |
|---|---|---|---|
| 1/f slope | LAI, ACC, MPFC, LAG, RAG | 39 | Dominant selected family across five- and ten-feature panels. Its recurrence suggests a weak aperiodic spectral-tilt contribution to CON-PSP separation, but the low classifier performance indicates poor individual-level coherence. |
| Beta peak frequency | ACC, LAI, RAI, LAG, MPFC | 17 | Appears only in ten-feature panels. It describes subtle shifts in beta-peak location rather than a strong beta-power marker, and should be interpreted as distributed weak beta-organisation information. |
| Relative gamma power | LAG | 4 | Selected mainly in effect-size and combined ten-feature panels. It contributes to the best D5 operating point through LAG, but the effect is weak and not a standalone marker. |

**Table 2.5. Full PSD classifier metrics for controls-PSP.**

| Panel | Model | Accuracy | Sensitivity | Specificity | Precision | F1 | MCC | AUC |
|---|---|---|---|---|---|---|---|---|
| P5 | Lin-SVM | 0.550 | 0.236 | 0.784 | 0.448 | 0.310 | 0.024 | 0.504 |
| P5 | RBF-SVM | 0.519 | 0.327 | 0.662 | 0.419 | 0.367 | -0.011 | 0.533 |
| P5 | RF | 0.535 | 0.455 | 0.595 | 0.455 | 0.455 | 0.049 | 0.522 |
| P5 | AdaBoost | 0.473 | 0.400 | 0.527 | 0.386 | 0.393 | -0.073 | 0.504 |
| P10 | Lin-SVM | 0.581 | 0.255 | 0.824 | 0.519 | 0.341 | 0.096 | 0.524 |

| Panel | Model | Accuracy | Sensitivity | Specificity | Precision | F1 | MCC | AUC |
|---|---|---|---|---|---|---|---|---|
| P10 | RBF-SVM | 0.434 | 0.618 | 0.297 | 0.395 | 0.482 | -0.089 | 0.460 |
| P10 | RF | 0.434 | 0.327 | 0.514 | 0.333 | 0.330 | -0.160 | 0.423 |
| P10 | AdaBoost | 0.473 | 0.436 | 0.500 | 0.393 | 0.414 | -0.063 | 0.442 |
| D5 | Lin-SVM | 0.566 | 0.127 | 0.892 | 0.467 | 0.200 | 0.030 | 0.480 |
| D5 | RBF-SVM | 0.496 | 0.273 | 0.662 | 0.375 | 0.316 | -0.070 | 0.453 |
| D5 | RF | 0.558 | 0.400 | 0.676 | 0.478 | 0.436 | 0.078 | 0.533 |
| D5 | AdaBoost | 0.589 | 0.509 | 0.649 | 0.519 | 0.514 | 0.158 | 0.582 |
| D10 | Lin-SVM | 0.574 | 0.200 | 0.851 | 0.500 | 0.286 | 0.068 | 0.538 |
| D10 | RBF-SVM | 0.481 | 0.582 | 0.405 | 0.421 | 0.489 | -0.013 | 0.471 |
| D10 | RF | 0.473 | 0.345 | 0.568 | 0.373 | 0.358 | -0.088 | 0.418 |
| D10 | AdaBoost | 0.450 | 0.345 | 0.527 | 0.352 | 0.349 | -0.128 | 0.437 |
| C5 | Lin-SVM | 0.550 | 0.236 | 0.784 | 0.448 | 0.310 | 0.024 | 0.504 |
| C5 | RBF-SVM | 0.519 | 0.327 | 0.662 | 0.419 | 0.367 | -0.011 | 0.533 |
| C5 | RF | 0.535 | 0.455 | 0.595 | 0.455 | 0.455 | 0.049 | 0.522 |
| C5 | AdaBoost | 0.473 | 0.400 | 0.527 | 0.386 | 0.393 | -0.073 | 0.504 |
| C10 | Lin-SVM | 0.574 | 0.200 | 0.851 | 0.500 | 0.286 | 0.068 | 0.538 |
| C10 | RBF-SVM | 0.481 | 0.582 | 0.405 | 0.421 | 0.489 | -0.013 | 0.471 |
| C10 | RF | 0.481 | 0.345 | 0.581 | 0.380 | 0.362 | -0.075 | 0.418 |
| C10 | AdaBoost | 0.473 | 0.364 | 0.554 | 0.377 | 0.370 | -0.083 | 0.427 |
| S5 | Lin-SVM | 0.550 | 0.236 | 0.784 | 0.448 | 0.310 | 0.024 | 0.504 |
| S5 | RBF-SVM | 0.519 | 0.327 | 0.662 | 0.419 | 0.367 | -0.011 | 0.533 |
| S5 | RF | 0.535 | 0.455 | 0.595 | 0.455 | 0.455 | 0.049 | 0.522 |
| S5 | AdaBoost | 0.473 | 0.400 | 0.527 | 0.386 | 0.393 | -0.073 | 0.504 |
| S10 | Lin-SVM | 0.574 | 0.200 | 0.851 | 0.500 | 0.286 | 0.068 | 0.538 |
| S10 | RBF-SVM | 0.481 | 0.582 | 0.405 | 0.421 | 0.489 | -0.013 | 0.471 |
| S10 | RF | 0.488 | 0.345 | 0.595 | 0.388 | 0.365 | -0.061 | 0.424 |
| S10 | AdaBoost | 0.481 | 0.382 | 0.554 | 0.389 | 0.385 | -0.064 | 0.454 |

**Table 2.6. PSD confusion-matrix counts for controls-PSP.**

| Panel | Model | TP | TN | FP | FN | N evaluated |
|---|---|---|---|---|---|---|
| P5 | Lin-SVM | 13 | 58 | 16 | 42 | 129 |
| P5 | RBF-SVM | 18 | 49 | 25 | 37 | 129 |
| P5 | RF | 25 | 44 | 30 | 30 | 129 |
| P5 | AdaBoost | 22 | 39 | 35 | 33 | 129 |
| P10 | Lin-SVM | 14 | 61 | 13 | 41 | 129 |
| P10 | RBF-SVM | 34 | 22 | 52 | 21 | 129 |
| P10 | RF | 18 | 38 | 36 | 37 | 129 |
| P10 | AdaBoost | 24 | 37 | 37 | 31 | 129 |
| D5 | Lin-SVM | 7 | 66 | 8 | 48 | 129 |
| D5 | RBF-SVM | 15 | 49 | 25 | 40 | 129 |
| D5 | RF | 22 | 50 | 24 | 33 | 129 |
| D5 | AdaBoost | 28 | 48 | 26 | 27 | 129 |
| D10 | Lin-SVM | 11 | 63 | 11 | 44 | 129 |
| D10 | RBF-SVM | 32 | 30 | 44 | 23 | 129 |
| D10 | RF | 19 | 42 | 32 | 36 | 129 |
| D10 | AdaBoost | 19 | 39 | 35 | 36 | 129 |
| C5 | Lin-SVM | 13 | 58 | 16 | 42 | 129 |
| C5 | RBF-SVM | 18 | 49 | 25 | 37 | 129 |
| C5 | RF | 25 | 44 | 30 | 30 | 129 |
| C5 | AdaBoost | 22 | 39 | 35 | 33 | 129 |
| C10 | Lin-SVM | 11 | 63 | 11 | 44 | 129 |
| C10 | RBF-SVM | 32 | 30 | 44 | 23 | 129 |
| C10 | RF | 19 | 43 | 31 | 36 | 129 |
| C10 | AdaBoost | 20 | 41 | 33 | 35 | 129 |
| S5 | Lin-SVM | 13 | 58 | 16 | 42 | 129 |
| S5 | RBF-SVM | 18 | 49 | 25 | 37 | 129 |
| S5 | RF | 25 | 44 | 30 | 30 | 129 |
| S5 | AdaBoost | 22 | 39 | 35 | 33 | 129 |

| Panel | Model | TP | TN | FP | FN | N evaluated |
|---|---|---|---|---|---|---|
| S10 | Lin-SVM | 11 | 63 | 11 | 44 | 129 |
| S10 | RBF-SVM | 32 | 30 | 44 | 23 | 129 |
| S10 | RF | 19 | 44 | 30 | 36 | 129 |
| S10 | AdaBoost | 21 | 41 | 33 | 34 | 129 |

##### *DCM-parameter classification*

DCM information gives stronger individual-level coherence than PSD. The best reported CON-versus-PSP DCM classifier was a linear SVM using the P10 panel, with accuracy 0.698, MCC 0.371 and AUC 0.770. This should be interpreted as moderate group-informed coherence in latent physiological space rather than prospective diagnostic performance. The current DCM statistics also show no FDR-surviving single parameter, so the result is best interpreted as a distributed timing/gain/coupling pattern rather than an isolated DCM marker.

**Table 2.7. DCM classification result for controls-PSP.**

| Contrast | Domain | Best model / panel | Accuracy | Specificity | MCC | AUC | Interpretation |
|---|---|---|---|---|---|---|---|
| CON vs PSP | DCM | Linear SVM, P10 | 0.698 | not supplied | 0.371 | 0.770 | Moderate DCM coherence and stronger than PSD; interpreted as a distributed latent timing/gain/coupling signal. |

**Table 2.8. Univariate DCM statistical context for controls-PSP.**

| Contrast | Valid tests | p < 0.05 | p < 0.01 | q < 0.05 | Min p | Min q | Dominant top-ranked families |
|---|---|---|---|---|---|---|---|
| CON vs PSP | 337 valid / 525 parameters | 21 | 1 | 0 | 0.00685 | 0.667 | H (3); AN{2} (2); A{1}, AN{1}, CV, a, c (1 each) |

**Table 2.9. Selected P10 DCM parameter panel for controls-PSP.**

| Order | Parameter | Family / location | Interpretation | Higher mean | Mean CON | Mean PSP | p | q | d |
|---|---|---|---|---|---|---|---|---|---|
| 1 | H(1,3,5) | PCC intrinsic coupling | Inhibitory-to-spiny-excitatory coupling in PCC. | CON | 0.080 | -0.009 | 0.00685 | 0.667 | 0.486 |
| 2 | A{1}(4,3) | Forward AMPA | RAI to MPFC AMPA-mediated effective coupling. | PSP | 0.136 | 0.409 | 0.016 | 0.667 | -0.465 |
| 3 | H(2,2,4) | MPFC self-coupling | Superficial-pyramidal self-coupling/gain in MPFC. | CON | 0.047 | -0.122 | 0.018 | 0.667 | 0.460 |
| 4 | c(1,5) | Specific sensor noise | Specific sensor-noise term; not direct synaptic physiology. | CON | -0.030 | -0.048 | 0.0203 | 0.667 | 0.443 |
| 5 | a(2,4) | Endogenous fluctuation | Modelled endogenous fluctuation term; not direct synaptic physiology. | PSP | 0.237 | 0.424 | 0.0224 | 0.667 | -0.440 |
| 6 | H(2,3,1) | LAI intrinsic coupling | Inhibitory-to-superficial-pyramidal coupling in LAI. | CON | 0.032 | 0.000 | 0.0284 | 0.667 | 0.380 |
| 7 | CV(2) | Superficial-pyramidal capacitance | Membrane capacitance of the superficial-pyramidal population. | PSP | 0.152 | 0.327 | 0.0308 | 0.667 | -0.404 |
| 8 | AN{2}(1,4) | Backward NMDA | MPFC to LAI NMDA-mediated backward coupling. | CON | 0.053 | -0.017 | 0.0325 | 0.667 | 0.364 |
| 9 | AN{2}(4,6) | Backward NMDA | LAG to MPFC NMDA-mediated backward coupling. | CON | 0.022 | -0.015 | 0.0331 | 0.667 | 0.352 |
| 10 | AN{1}(7,2) | Forward NMDA | ACC to RAG NMDA-mediated forward coupling. | PSP | -0.020 | 0.035 | 0.0353 | 0.667 | -0.417 |

##### *Contrast-level interpretation*

Controls-PSP shows a clearer separation between clinical/DCM and empirical PSD readouts. The PSD classification is weak, with the best model reaching only accuracy 0.589, MCC 0.158 and AUC 0.582, and the selected features mainly reflect widespread aperiodic-slope trends plus limited beta/gamma contributions. Together with the focal LAI/RAG low-frequency bin wise effects and absence of FDR-surviving PSD features, this supports a weak and distributed empirical spectral phenotype rather than a robust individual-level PSD marker. The DCM result is stronger but still moderate, with linear SVM P10 accuracy 0.698, MCC 0.371 and AUC 0.770. The overall interpretation is therefore that PSP-control physiology is more coherently expressed in latent DCM timing/gain/coupling space than in empirical PSD-feature space.

#### 3. Controls versus pooled patients

##### *PSD-feature classification*

The pooled-patient analysis used 174 participants: 74 controls and 100 pooled patients (45 bvFTD and 55 PSP). The leading audited threshold-independent result used an RBF-SVM with a compact five-feature beta-residual/theta panel, reaching accuracy 0.586, sensitivity 0.660, specificity 0.486, precision 0.635, F1-score 0.647, MCC 0.148 and AUC 0.591 (TP 66, TN 36, FP 38, FN 34; N = 174). A sensitivity-biased AdaBoost solution on the same five-feature panel gave higher apparent accuracy (0.621) and sensitivity (0.880), but low specificity (0.270), MCC 0.162 and AUC 0.476, and was therefore not treated as stronger physiological evidence.

The selected PSD features were dominated by beta residual maximum in RAI, RAG, MPFC and ACC, together with relative theta power in RAI. This agrees with the univariate PSD feature report: no derived PSD feature survived BH-FDR correction in the pooled contrast, and the leading uncorrected effects were beta residual maxima across most sources, RAI relative theta, and posterior alpha residual maxima. The interpretation is a weak transdiagnostic spectral axis rather than a robust individual-level PSD marker.

**Table 3.1. PSD classification design for control-pooled patients.**

| Item | Value |
|---|---|
| Contrast | Controls versus pooled patients (bvFTD + PSP) |
| Sample size | N = 174 total; controls = 74; pooled patients = 100 |
| Pooled-patient composition | 45 bvFTD + 55 PSP |
| Label coding | Controls = 0; pooled patients = 1 |
| Positive class for sensitivity / precision / F1 | Pooled patients |
| Negative class for specificity | Controls |
| Evaluation scheme | Leave-one-out cross-validation |
| Classifier families | Linear SVM, RBF-SVM, Random Forest, AdaBoost |
| Feature panels | P5/P10, D5/D10, C5/C10, S5/S10 |

**Table 3.2. PSD operating points for controls -pooled patients.**

| Summary criterion | Panel / model | Performance metrics | Confusion matrix | Interpretation |
|---|---|---|---|---|
| Best threshold-independent PSD result | Compact five-feature beta-residual/theta panel / RBF-SVM | Acc 0.586; Sens 0.660; Spec 0.486; Prec 0.635; F1 0.647; MCC 0.148; AUC 0.591 | TP 66; TN 36; FP 38; FN 34; N 174 | Weak. |
| Sensitivity-biased higher-accuracy solution | Same compact five-feature panel / AdaBoost | Acc 0.621; Sens 0.880; Spec 0.270; F1 0.722; MCC 0.162; AUC 0.476 | AUC and specificity were poor. | High sensitivity mainly reflects patient-biased assignment. |
| Statistical context | PSD feature statistics | 105 features; 5 uncorrected p<0.05; 0 p<0.01; 0 q<0.05; min p 0.029; min q 0.446 | Not applicable | No corrected pooled-patient PSD feature survived FDR. |

**Table 3.3. PSD feature families supporting pooled-patient interpretation.**

| Feature family | Sources / features | Direction and interpretation |
|---|---|---|
| Beta residual maximum | RAI, RAG, MPFC, ACC, LAG, PCC, LAI | Controls showed higher beta-residual structure than pooled patients. This supports a weak common disease axis involving reduced organised beta rhythmicity. |
| Relative theta power | RAI | Pooled patients showed higher RAI theta, adding a weak slowing-related component. |
| Alpha residual maximum | LAG and PCC | Controls showed higher posterior alpha residual structure; this contributes only weakly to the transdiagnostic signal. |

##### *DCM-parameter classification*

DCM prediction was modestly stronger than PSD in the pooled contrast. The strongest DCM classification operating point results achieved with Random Forest using the P10/STABLE_TOP10 ten-parameter panel: accuracy 0.632, sensitivity 0.720, specificity 0.514, precision 0.667, F1-score 0.692, MCC 0.238 and AUC 0.639 (TP 72, TN 38, FP 36, FN 28). The highest threshold-independent AUC was obtained by a linear SVM using the compact five-parameter panel [CV(2), H(2,2,4), A{1}(4,3), H(2,2,2), A{1}(5,2)] (AUC 0.648; accuracy 0.598; MCC 0.174).

Twenty-four DCM parameters reached uncorrected $p < 0.05$ and six reached $p < 0.01$, but no parameter survived BH-FDR correction (minimum $p = 0.00282$; minimum $q = 0.410$). The classification result is therefore best interpreted as weak-to-modest multivariate coherence in latent physiological space, not as a corrected single-parameter biomarker.

**Table 3.4. DCM feature-panel definitions for control-pooled patients.**

| Panel | Selection principle | Panel size | Interpretation |
|---|---|---|---|
| P5 / P10 | Smallest uncorrected Welch p-values | 5 / 10 | Prioritises statistically prominent DCM parameters. |
| D5 / D10 | Largest absolute Cohen d values | 5 / 10 | Prioritises parameters with larger group separation. |
| C5 / C10 | Combined p-value and effect-size ranking | 5 / 10 | Balances strength of evidence and effect magnitude. |
| S5 / S10 | Combined score plus recurrence/stability bonus | 5 / 10 | Favours parameters recurring across disease contrasts. |

**Table 3.5. Selected DCM parameter panels for controls-pooled patients.**

| Panel | Selected DCM parameters |
|---|---|
| P5 | 1. CV(2); 2. H(2,2,4); 3. A{1}(4,3); 4. H(2,2,2); 5. A{1}(5,2) |
| P10 | 1. CV(2); 2. H(2,2,4); 3. A{1}(4,3); 4. H(2,2,2); 5. A{1}(5,2); 6. A{1}(4,5); 7. H(1,1,2); 8. A{1}(3,7); 9. T(1,2); 10. c(1,5) |
| D5 | 1. CV(2); 2. H(2,2,4); 3. A{1}(4,3); 4. H(2,2,2); 5. A{1}(5,2) |
| D10 | 1. CV(2); 2. H(2,2,4); 3. A{1}(4,3); 4. H(2,2,2); 5. A{1}(5,2); 6. A{1}(4,5); 7. H(1,1,2); 8. T(1,2); 9. c(1,5); 10. A{1}(3,7) |
| C5 | 1. CV(2); 2. H(2,2,4); 3. A{1}(4,3); 4. H(2,2,2); 5. A{1}(5,2) |
| C10 | 1. CV(2); 2. H(2,2,4); 3. A{1}(4,3); 4. H(2,2,2); 5. A{1}(5,2); 6. A{1}(4,5); 7. H(1,1,2); 8. T(1,2); 9. A{1}(3,7); 10. c(1,5) |
| S5 | 1. CV(2); 2. H(2,2,4); 3. A{1}(5,2); 4. A{1}(4,3); 5. H(2,2,2) |
| S10 | 1. CV(2); 2. H(2,2,4); 3. A{1}(5,2); 4. A{1}(4,3); 5. H(2,2,2); 6. A{1}(4,5); 7. H(1,1,2); 8. T(1,2); 9. A{1}(3,7); 10. c(1,5) |

**Table 3.6. Physiological interpretation of selected DCM parameters in the pooled contrast.**

| Parameter | Meaning in this model | Interpretive role in this contrast |
|---|---|---|
| CV(2) | Membrane capacitance of the superficial pyramidal population. | Recurrent core term; suggests that the common disease-control axis partly involves superficial-pyramidal excitability/capacitance. |
| H(2,2,4) | Superficial-pyramidal self-coupling in MPFC. | Local gain/self-coupling term within a frontal default-mode/salience-interacting node. |
| A{1}(4,3) | Forward AMPA coupling from RAI to MPFC. | Right-anterior-insular to medial-prefrontal communication; part of the shared AMPA-mediated patient-control axis. |
| H(2,2,2) | Superficial-pyramidal self-coupling in ACC. | Local cingulate gain term recurring across patient-control panels. |
| A{1}(5,2) | Forward AMPA coupling from ACC to PCC. | Links salience/cingulate dynamics to posterior default-mode cortex. |
| A{1}(4,5) | Forward AMPA coupling from PCC to MPFC. | Posterior-to-medial-prefrontal communication within the default-mode component. |
| H(1,1,2) | Spiny-excitatory self-coupling in ACC. | Additional ACC intrinsic coupling term. |
| A{1}(3,7) | Forward AMPA coupling from RAG to RAI. | Posterior angular-gyrus to right-anterior-insular pathway. |
| T(1,2) | GABA_A inhibitory time constant in LAI. | Inhibitory timing term included in ten-parameter patient-control panels. |

| Parameter | Meaning in this model | Interpretive role in this contrast |
|---|---|---|
| c(1,5) | Specific sensor-noise component. | Modelled observation/noise term; should not be interpreted as direct synaptic physiology. |

**Table 3.7. Best DCM classifier within each feature panel for controls-pooled patients.**

| Panel | Best AUC model | AUC | Accuracy | Sensitivity | Specificity | F1 | MCC | Best operating model | Operating-point metrics |
|---|---|---|---|---|---|---|---|---|---|
| P5 | Lin-SVM | 0.648 | 0.598 | 0.660 | 0.514 | 0.653 | 0.174 | RBF-SVM | Acc 0.621; F1 0.694; MCC 0.205 |
| P10 | RF | 0.639 | 0.632 | 0.720 | 0.514 | 0.692 | 0.238 | RF | Acc 0.632; F1 0.692; MCC 0.238 |
| D5 | Lin-SVM | 0.648 | 0.598 | 0.660 | 0.514 | 0.653 | 0.174 | RBF-SVM | Acc 0.621; F1 0.694; MCC 0.205 |
| D10 | RF | 0.638 | 0.626 | 0.710 | 0.514 | 0.686 | 0.227 | RF | Acc 0.626; F1 0.686; MCC 0.227 |
| C5 | Lin-SVM | 0.648 | 0.598 | 0.660 | 0.514 | 0.653 | 0.174 | RBF-SVM | Acc 0.621; F1 0.694; MCC 0.205 |
| C10 | RF | 0.638 | 0.626 | 0.710 | 0.514 | 0.686 | 0.227 | RF | Acc 0.626; F1 0.686; MCC 0.227 |
| S5 | Lin-SVM | 0.648 | 0.598 | 0.660 | 0.514 | 0.653 | 0.174 | RBF-SVM | Acc 0.621; F1 0.694; MCC 0.205 |
| S10 | RF | 0.639 | 0.632 | 0.720 | 0.514 | 0.692 | 0.238 | RF | Acc 0.632; F1 0.692; MCC 0.238 |

**Table 3.8. DCM classifier metrics for controls-patients.**

| Panel | Model | N | Acc | Sens | Spec | Prec | F1 | MCC | AUC | TP | TN | FP | FN |
|---|---|---|---|---|---|---|---|---|---|---|---|---|---|
| P5 | Lin-SVM | 174 | 0.598 | 0.660 | 0.514 | 0.647 | 0.653 | 0.174 | 0.648 | 66 | 38 | 36 | 34 |
| P5 | RBF-SVM | 174 | 0.621 | 0.750 | 0.446 | 0.647 | 0.694 | 0.205 | 0.598 | 75 | 33 | 41 | 25 |
| P5 | RF | 174 | 0.552 | 0.650 | 0.419 | 0.602 | 0.625 | 0.070 | 0.568 | 65 | 31 | 43 | 35 |
| P5 | AdaBoost | 174 | 0.569 | 0.650 | 0.459 | 0.619 | 0.634 | 0.111 | 0.544 | 65 | 34 | 40 | 35 |
| P10 | Lin-SVM | 174 | 0.592 | 0.640 | 0.527 | 0.646 | 0.643 | 0.167 | 0.593 | 64 | 39 | 35 | 36 |
| P10 | RBF-SVM | 174 | 0.563 | 0.700 | 0.378 | 0.603 | 0.648 | 0.082 | 0.567 | 70 | 28 | 46 | 30 |
| P10 | RF | 174 | 0.632 | 0.720 | 0.514 | 0.667 | 0.692 | 0.238 | 0.639 | 72 | 38 | 36 | 28 |
| P10 | AdaBoost | 174 | 0.603 | 0.650 | 0.541 | 0.657 | 0.653 | 0.190 | 0.630 | 65 | 40 | 34 | 35 |
| D5 | Lin-SVM | 174 | 0.598 | 0.660 | 0.514 | 0.647 | 0.653 | 0.174 | 0.648 | 66 | 38 | 36 | 34 |
| D5 | RBF-SVM | 174 | 0.621 | 0.750 | 0.446 | 0.647 | 0.694 | 0.205 | 0.598 | 75 | 33 | 41 | 25 |
| D5 | RF | 174 | 0.552 | 0.650 | 0.419 | 0.602 | 0.625 | 0.070 | 0.568 | 65 | 31 | 43 | 35 |
| D5 | AdaBoost | 174 | 0.569 | 0.650 | 0.459 | 0.619 | 0.634 | 0.111 | 0.544 | 65 | 34 | 40 | 35 |
| D10 | Lin-SVM | 174 | 0.592 | 0.640 | 0.527 | 0.646 | 0.643 | 0.167 | 0.593 | 64 | 39 | 35 | 36 |
| D10 | RBF-SVM | 174 | 0.563 | 0.700 | 0.378 | 0.603 | 0.648 | 0.082 | 0.567 | 70 | 28 | 46 | 30 |
| D10 | RF | 174 | 0.626 | 0.710 | 0.514 | 0.664 | 0.686 | 0.227 | 0.638 | 71 | 38 | 36 | 29 |
| D10 | AdaBoost | 174 | 0.609 | 0.660 | 0.541 | 0.660 | 0.660 | 0.201 | 0.629 | 66 | 40 | 34 | 34 |
| C5 | Lin-SVM | 174 | 0.598 | 0.660 | 0.514 | 0.647 | 0.653 | 0.174 | 0.648 | 66 | 38 | 36 | 34 |
| C5 | RBF-SVM | 174 | 0.621 | 0.750 | 0.446 | 0.647 | 0.694 | 0.205 | 0.598 | 75 | 33 | 41 | 25 |
| C5 | RF | 174 | 0.552 | 0.650 | 0.419 | 0.602 | 0.625 | 0.070 | 0.568 | 65 | 31 | 43 | 35 |
| C5 | AdaBoost | 174 | 0.569 | 0.650 | 0.459 | 0.619 | 0.634 | 0.111 | 0.544 | 65 | 34 | 40 | 35 |
| C10 | Lin-SVM | 174 | 0.592 | 0.640 | 0.527 | 0.646 | 0.643 | 0.167 | 0.593 | 64 | 39 | 35 | 36 |
| C10 | RBF-SVM | 174 | 0.563 | 0.700 | 0.378 | 0.603 | 0.648 | 0.082 | 0.567 | 70 | 28 | 46 | 30 |
| C10 | RF | 174 | 0.626 | 0.710 | 0.514 | 0.664 | 0.686 | 0.227 | 0.638 | 71 | 38 | 36 | 29 |
| C10 | AdaBoost | 174 | 0.603 | 0.650 | 0.541 | 0.657 | 0.653 | 0.190 | 0.629 | 65 | 40 | 34 | 35 |
| S5 | Lin-SVM | 174 | 0.598 | 0.660 | 0.514 | 0.647 | 0.653 | 0.174 | 0.648 | 66 | 38 | 36 | 34 |
| S5 | RBF-SVM | 174 | 0.621 | 0.750 | 0.446 | 0.647 | 0.694 | 0.205 | 0.598 | 75 | 33 | 41 | 25 |
| S5 | RF | 174 | 0.563 | 0.670 | 0.419 | 0.609 | 0.638 | 0.091 | 0.567 | 67 | 31 | 43 | 33 |
| S5 | AdaBoost | 174 | 0.529 | 0.610 | 0.419 | 0.587 | 0.598 | 0.029 | 0.544 | 61 | 31 | 43 | 39 |
| S10 | Lin-SVM | 174 | 0.592 | 0.640 | 0.527 | 0.646 | 0.643 | 0.167 | 0.593 | 64 | 39 | 35 | 36 |
| S10 | RBF-SVM | 174 | 0.563 | 0.700 | 0.378 | 0.603 | 0.648 | 0.082 | 0.567 | 70 | 28 | 46 | 30 |
| S10 | RF | 174 | 0.632 | 0.720 | 0.514 | 0.667 | 0.692 | 0.238 | 0.639 | 72 | 38 | 36 | 28 |
| S10 | AdaBoost | 174 | 0.586 | 0.650 | 0.500 | 0.637 | 0.644 | 0.151 | 0.623 | 65 | 37 | 37 | 35 |

**Table 3.9. Top uncorrected DCM parameter effects for controls-pooled patients**

| Rank | Parameter | Higher mean | Mean A | Mean B | p_Welch | q_BH | Cohen d |
|---|---|---|---|---|---|---|---|
| 1 | CV(2) | Patient | 0.152 | 0.348 | 0.003 | 0.410 | -0.452 |
| 2 | H(2,2,4) | CON | 0.047 | -0.114 | 0.003 | 0.410 | 0.432 |
| 3 | A{1}(4,3) | Patient | 0.136 | 0.388 | 0.005 | 0.410 | -0.406 |
| 4 | H(2,2,2) | CON | 0.126 | 0.015 | 0.006 | 0.410 | 0.405 |
| 5 | A{1}(5,2) | Patient | -0.004 | 0.070 | 0.006 | 0.410 | -0.403 |
| 6 | A{1}(4,5) | Patient | -0.068 | 0.046 | 0.009 | 0.497 | -0.371 |
| 7 | H(1,1,2) | CON | -0.008 | -0.098 | 0.012 | 0.535 | 0.367 |
| 8 | A{1}(3,7) | Patient | 0.011 | 0.150 | 0.016 | 0.535 | -0.346 |
| 9 | T(1,2) | Patient | 0.078 | 0.146 | 0.016 | 0.535 | -0.357 |
| 10 | c(1,5) | CON | -0.030 | -0.044 | 0.017 | 0.535 | 0.353 |

Controls- pooled patients produce weaker PSD and DCM individual-level coherence than the syndrome-specific contrasts. PSD classification is dominated by a shallow reduction in beta residual structure and weak slowing-related features. DCM classification is slightly stronger but remains limited, with a recurrent gain/coupling core and no FDR-surviving single DCM parameter. Pooling preserves a common disease-control axis but reduces syndrome-specific physiological contrast.

#### 4. bvFTD versus PSP

##### *PSD-feature classification*

The selected PSD panels were dominated by 1/f slope across the SN-DMN sources, with relative beta contributions in RAI, LAI and RAG. This supports an aperiodic/beta syndrome rather than diagnostic.

**Table 4.1. PSD classification design and label coding for bvFTD - PSP.**

| Item | Value |
|---|---|
| Contrast | bvFTD versus PSP |
| Sample size | N = 100 total; bvFTD = 45; PSP = 55 |
| Label coding | PSP = 0; bvFTD = 1 |
| Positive class for sensitivity / precision / F1 | bvFTD |
| Negative class for specificity | PSP |
| Evaluation scheme | Leave-one-out cross-validation |
| Classifier families | Linear SVM, RBF-SVM, Random Forest, AdaBoost |
| Feature panels | P5/P10, D5/D10, C5/C10, S5/S10 |

**Table 4.2.PSD feature-panel and classification summary for bvFTD- PSP.**

| Panel | Selected PSD feature pattern | Best AUC model | AUC | Acc. | Sens. | Spec. | MCC | Best operating point |
|---|---|---|---|---|---|---|---|---|
| P5 | 1/f slope: ACC, LAG, LAI, MPFC, RAG | Linear SVM | 0.635 | 0.590 | 0.489 | 0.673 | 0.164 | Random forest: Acc 0.630; F1 0.575; MCC 0.248 |
| P10 | 1/f slope: LAI, ACC, RAI, MPFC, PCC, LAG, RAG; RelBeta: RAI, LAI, RAG | Linear SVM | 0.652 | 0.610 | 0.533 | 0.673 | 0.208 | Random forest: Acc 0.620; F1 0.568; MCC 0.229 |
| D5 | 1/f slope: ACC, LAG, MPFC, LAI, RAG | Random forest | 0.639 | 0.620 | 0.533 | 0.691 | 0.227 | AdaBoost: Acc 0.640; F1 0.571; MCC 0.266 |
| D10 | 1/f slope: LAI, ACC, RAI, MPFC, PCC, LAG, RAG; RelBeta: RAI, LAI, RAG | Linear SVM | 0.652 | 0.610 | 0.533 | 0.673 | 0.208 | AdaBoost: Acc 0.630; F1 0.593; MCC 0.254 |
| C5 | 1/f slope: ACC, LAG, LAI, MPFC, RAG | Linear SVM | 0.635 | 0.590 | 0.489 | 0.673 | 0.164 | Random forest: Acc 0.630; F1 0.575; MCC 0.248 |
| C10 | 1/f slope: LAI, ACC, RAI, MPFC, PCC, LAG, RAG; RelBeta: RAI, LAI, RAG | Linear SVM | 0.652 | 0.610 | 0.533 | 0.673 | 0.208 | Random forest: Acc 0.620; F1 0.568; MCC 0.229 |
| S5 | 1/f slope: ACC, LAG, LAI, MPFC, RAG | Linear SVM | 0.635 | 0.590 | 0.489 | 0.673 | 0.164 | Random forest: Acc 0.630; F1 0.575; MCC 0.248 |
| S10 | 1/f slope: LAI, ACC, RAI, MPFC, PCC, LAG, RAG; RelBeta: RAI, LAI, RAG | Linear SVM | 0.652 | 0.610 | 0.533 | 0.673 | 0.208 | Random forest: Acc 0.620; F1 0.568; MCC 0.229 |

##### *DCM-parameter classification*

The DCM classification best operating point was a linear SVM using a compact five-parameter DCM panel. The same linear solution appeared for the P5, D5, C5 and S5 panels, with accuracy 0.740, sensitivity 0.800, specificity 0.667, precision 0.746, F1-score 0.772, MCC 0.472 and AUC 0.798. The selected parameters were T(7,1), AN{1}(4,2), A{1}(7,6), AN{2}(1,4) and D(2). This supports a compact timing/coupling syndrome axis involving posterior AMPA timing, NMDA-mediated salience/default-mode coupling, angular-gyrus AMPA communication and extrinsic delay.

**Table 4.3. Best DCM classification result for bvFTD-PSP.**

| Classifier | Panel | Selected DCM parameters | Accuracy | Sensitivity | Specificity | Precision | F1 | MCC | AUC | TP | TN | FP | FN |
|---|---|---|---|---|---|---|---|---|---|---|---|---|---|
| Linear SVM | P5/D5/C5/S5 compact five-parameter panel | T(7,1), AN{1}(4,2), A{1}(7,6), AN{2}(1,4), D(2) | 0.740 | 0.800 | 0.667 | 0.746 | 0.772 | 0.472 | 0.798 | 44 | 30 | 15 | 11 |

**Table 4.4. DCM selected-parameter panels for bvFTD-PSP.**

| Panel | n | Selected DCM parameters |
|---|---|---|
| P5 | 5 | T(7,1); AN{1}(4,2); A{1}(7,6); D(2); AN{2}(1,4) |
| P10 | 10 | T(7,1); AN{1}(4,2); A{1}(7,6); D(2); AN{2}(1,4); d(2,7); AN{2}(5,7); L(5); AN{2}(7,6); d(3,7) |
| D5 | 5 | AN{1}(4,2); T(7,1); AN{2}(1,4); A{1}(7,6); D(2) |
| D10 | 10 | AN{1}(4,2); T(7,1); AN{2}(1,4); A{1}(7,6); D(2); L(5); d(2,7); AN{2}(7,6); d(3,7); AN{2}(5,7) |
| C5 | 5 | T(7,1); AN{1}(4,2); A{1}(7,6); AN{2}(1,4); D(2) |
| C10 | 10 | T(7,1); AN{1}(4,2); A{1}(7,6); AN{2}(1,4); D(2); d(2,7); L(5); AN{2}(5,7); AN{2}(7,6); d(3,7) |
| S5 | 5 | T(7,1); AN{1}(4,2); AN{2}(1,4); A{1}(7,6); D(2) |
| S10 | 10 | T(7,1); AN{1}(4,2); AN{2}(1,4); A{1}(7,6); D(2); d(2,7); L(5); AN{2}(7,6); AN{2}(5,7); d(3,7) |

##### Table 4.5. DCM classification report.

| Panel | Best AUC model | AUC | Accuracy | Sensitivity | Specificity | F1 | MCC | Best operating model | Operating-point metrics |
|---|---|---|---|---|---|---|---|---|---|
| P5 | Lin-SVM | 0.798 | 0.740 | 0.800 | 0.667 | 0.772 | 0.472 | Lin-SVM | Acc 0.740; F1 0.772; MCC 0.472 |
| P10 | Lin-SVM | 0.783 | 0.690 | 0.764 | 0.600 | 0.730 | 0.369 | Lin-SVM | Acc 0.690; F1 0.730; MCC 0.369 |
| D5 | Lin-SVM | 0.798 | 0.740 | 0.800 | 0.667 | 0.772 | 0.472 | Lin-SVM | Acc 0.740; F1 0.772; MCC 0.472 |
| D10 | Lin-SVM | 0.783 | 0.690 | 0.764 | 0.600 | 0.730 | 0.369 | Lin-SVM | Acc 0.690; F1 0.730; MCC 0.369 |
| C5 | Lin-SVM | 0.798 | 0.740 | 0.800 | 0.667 | 0.772 | 0.472 | Lin-SVM | Acc 0.740; F1 0.772; MCC 0.472 |
| C10 | Lin-SVM | 0.783 | 0.690 | 0.764 | 0.600 | 0.730 | 0.369 | Lin-SVM | Acc 0.690; F1 0.730; MCC 0.369 |
| S5 | Lin-SVM | 0.798 | 0.740 | 0.800 | 0.667 | 0.772 | 0.472 | Lin-SVM | Acc 0.740; F1 0.772; MCC 0.472 |
| S10 | Lin-SVM | 0.783 | 0.690 | 0.764 | 0.600 | 0.730 | 0.369 | Lin-SVM | Acc 0.690; F1 0.730; MCC 0.369 |

##### Table 4.6. DCM classifier metrics for bvFTD-PSP.

| Panel | Model | N | Acc. | Sens. | Spec. | Prec. | F1 | MCC | AUC | TP | TN | FP | FN |
|---|---|---|---|---|---|---|---|---|---|---|---|---|---|
| P5 | Lin-SVM | 100 | 0.740 | 0.800 | 0.667 | 0.746 | 0.772 | 0.472 | 0.798 | 44 | 30 | 15 | 11 |
| P5 | RBF-SVM | 100 | 0.540 | 0.709 | 0.333 | 0.565 | 0.629 | 0.046 | 0.579 | 39 | 15 | 30 | 16 |
| P5 | RF | 100 | 0.630 | 0.673 | 0.578 | 0.661 | 0.667 | 0.251 | 0.623 | 37 | 26 | 19 | 18 |
| P5 | AdaBoost | 100 | 0.600 | 0.691 | 0.489 | 0.623 | 0.655 | 0.183 | 0.600 | 38 | 22 | 23 | 17 |
| P10 | Lin-SVM | 100 | 0.690 | 0.764 | 0.600 | 0.700 | 0.730 | 0.369 | 0.783 | 42 | 27 | 18 | 13 |
| P10 | RBF-SVM | 100 | 0.610 | 0.764 | 0.422 | 0.618 | 0.683 | 0.198 | 0.605 | 42 | 19 | 26 | 13 |
| P10 | RF | 100 | 0.610 | 0.655 | 0.556 | 0.643 | 0.649 | 0.211 | 0.662 | 36 | 25 | 20 | 19 |
| P10 | AdaBoost | 100 | 0.520 | 0.945 | 0.000 | 0.536 | 0.684 | -0.159 | 0.615 | 52 | 0 | 45 | 3 |
| D5 | Lin-SVM | 100 | 0.740 | 0.800 | 0.667 | 0.746 | 0.772 | 0.472 | 0.798 | 44 | 30 | 15 | 11 |
| D5 | RBF-SVM | 100 | 0.540 | 0.709 | 0.333 | 0.565 | 0.629 | 0.046 | 0.579 | 39 | 15 | 30 | 16 |
| D5 | RF | 100 | 0.630 | 0.673 | 0.578 | 0.661 | 0.667 | 0.251 | 0.633 | 37 | 26 | 19 | 18 |
| D5 | AdaBoost | 100 | 0.640 | 0.764 | 0.489 | 0.646 | 0.700 | 0.263 | 0.544 | 42 | 22 | 23 | 13 |
| D10 | Lin-SVM | 100 | 0.690 | 0.764 | 0.600 | 0.700 | 0.730 | 0.369 | 0.783 | 42 | 27 | 18 | 13 |
| D10 | RBF-SVM | 100 | 0.610 | 0.764 | 0.422 | 0.618 | 0.683 | 0.198 | 0.605 | 42 | 19 | 26 | 13 |
| D10 | RF | 100 | 0.600 | 0.655 | 0.533 | 0.632 | 0.643 | 0.189 | 0.668 | 36 | 24 | 21 | 19 |
| D10 | AdaBoost | 100 | 0.520 | 0.945 | 0.000 | 0.536 | 0.684 | -0.159 | 0.621 | 52 | 0 | 45 | 3 |
| C5 | Lin-SVM | 100 | 0.740 | 0.800 | 0.667 | 0.746 | 0.772 | 0.472 | 0.798 | 44 | 30 | 15 | 11 |
| C5 | RBF-SVM | 100 | 0.540 | 0.709 | 0.333 | 0.565 | 0.629 | 0.046 | 0.579 | 39 | 15 | 30 | 16 |
| C5 | RF | 100 | 0.630 | 0.673 | 0.578 | 0.661 | 0.667 | 0.251 | 0.621 | 37 | 26 | 19 | 18 |
| C5 | AdaBoost | 100 | 0.610 | 0.709 | 0.489 | 0.629 | 0.667 | 0.203 | 0.592 | 39 | 22 | 23 | 16 |
| C10 | Lin-SVM | 100 | 0.690 | 0.764 | 0.600 | 0.700 | 0.730 | 0.369 | 0.783 | 42 | 27 | 18 | 13 |
| C10 | RBF-SVM | 100 | 0.610 | 0.764 | 0.422 | 0.618 | 0.683 | 0.198 | 0.605 | 42 | 19 | 26 | 13 |
| C10 | RF | 100 | 0.610 | 0.655 | 0.556 | 0.643 | 0.649 | 0.211 | 0.661 | 36 | 25 | 20 | 19 |
| C10 | AdaBoost | 100 | 0.520 | 0.945 | 0.000 | 0.536 | 0.684 | -0.159 | 0.615 | 52 | 0 | 45 | 3 |
| S5 | Lin-SVM | 100 | 0.740 | 0.800 | 0.667 | 0.746 | 0.772 | 0.472 | 0.798 | 44 | 30 | 15 | 11 |
| S5 | RBF-SVM | 100 | 0.540 | 0.709 | 0.333 | 0.565 | 0.629 | 0.046 | 0.579 | 39 | 15 | 30 | 16 |
| S5 | RF | 100 | 0.640 | 0.673 | 0.600 | 0.673 | 0.673 | 0.273 | 0.622 | 37 | 27 | 18 | 18 |
| S5 | AdaBoost | 100 | 0.590 | 0.709 | 0.444 | 0.609 | 0.655 | 0.159 | 0.594 | 39 | 20 | 25 | 16 |
| S10 | Lin-SVM | 100 | 0.690 | 0.764 | 0.600 | 0.700 | 0.730 | 0.369 | 0.783 | 42 | 27 | 18 | 13 |
| S10 | RBF-SVM | 100 | 0.610 | 0.764 | 0.422 | 0.618 | 0.683 | 0.198 | 0.605 | 42 | 19 | 26 | 13 |
| S10 | RF | 100 | 0.610 | 0.655 | 0.556 | 0.643 | 0.649 | 0.211 | 0.661 | 36 | 25 | 20 | 19 |
| S10 | AdaBoost | 100 | 0.520 | 0.945 | 0.000 | 0.536 | 0.684 | -0.159 | 0.621 | 52 | 0 | 45 | 3 |

##### Table 4.7. Interpretation of the compact five-parameter DCM panel selected for bvFTD-PSP.

| Parameter | Parameter meaning | Interpretation in this contrast |
|---|---|---|
| T(7,1) | RAG AMPA-mediated excitatory time constant. | Higher in PSP in the univariate DCM table. Its selection indicates that posterior association-network excitatory timing contributes to syndrome separation. |
| AN{1}(4,2) | Forward NMDA coupling from ACC to MPFC. | Higher in bvFTD. This supports involvement of slower receptor-specific integration from salience/cingulate cortex to medial prefrontal cortex. |
| A{1}(7,6) | Forward AMPA coupling from LAG to RAG. | Higher in PSP. This term contributes a posterior angular-gyrus AMPA communication component to the linear syndrome axis. |
| AN{2}(1,4) | Backward NMDA coupling from MPFC to LAI. | Higher in bvFTD. This links medial prefrontal/default-mode activity to anterior insula through slower NMDA-mediated feedback. |
| D(2) | Extrinsic conduction/transmission delay. | Higher in PSP. Its selection suggests that syndrome separation includes a distributed timing component, not only local gain or coupling. |

##### Table 4.8. Top uncorrected DCM parameter effects for bvFTD-PSP.

| Rank | Parameter | Higher mean | Mean A | Mean B | p_Welch | q_BH | Cohen's d |
|---|---|---|---|---|---|---|---|
| 1 | T(7,1) | PSP | -0.027 | 0.036 | 0.002 | 0.723 | -0.622 |
| 2 | AN{1}(4,2) | bvFTD | 0.148 | -0.055 | 0.005 | 0.781 | 0.634 |
| 3 | A{1}(7,6) | PSP | -0.031 | 0.166 | 0.009 | 0.797 | -0.526 |
| 4 | D(2) | PSP | 0.027 | 0.232 | 0.012 | 0.797 | -0.494 |
| 5 | AN{2}(1,4) | bvFTD | 0.143 | -0.017 | 0.013 | 0.797 | 0.556 |
| 6 | d(2,7) | bvFTD | 0.008 | -0.000 | 0.020 | 0.797 | 0.488 |
| 7 | AN{2}(5,7) | PSP | -0.006 | 0.012 | 0.021 | 0.797 | -0.446 |
| 8 | L(5) | bvFTD | 0.294 | 0.179 | 0.021 | 0.797 | 0.492 |
| 9 | AN{2}(7,6) | PSP | -0.063 | 0.041 | 0.023 | 0.797 | -0.454 |
| 10 | d(3,7) | bvFTD | 0.007 | -0.002 | 0.029 | 0.797 | 0.452 |

The bvFTD-PSP classification appendix supports a layered interpretation. PSD classification remains physiologically meaningful but modest, with a syndrome axis dominated by widespread 1/f slope and relative beta features. DCM classification shows stronger individual-level coherence, with a compact linear five-parameter solution reaching AUC 0.798 and MCC 0.472. The DCM result strengthens the conclusion that the direct syndrome difference is expressed through posterior timing, AMPA/NMDA-mediated effective connectivity, and extrinsic delay within the SN-DMN source space.

### Appendix D: The following abbreviations are used throughout the manuscript.

| Abbreviation | Full term / meaning | Use in this manuscript |
|---|---|---|
| ACE-R | Addenbrooke's Cognitive Examination-Revised | Global cognitive assessment is used as a clinical feature. |
| ACC | Anterior cingulate cortex | Salience-network source region. |
| AdaBoost | Adaptive Boosting | Ensemble classifier used to test threshold-like or interaction-based feature structure. |
| LAI | Left anterior insula | Salience-network source region. |
| AMPA | alpha-amino-3-hydroxy-5-methyl-4-isoxazolepropionic acid receptor | Fast excitatory receptor class represented in DCM coupling terms. |
| AUC | Area under the receiver operating characteristic curve | Threshold-independent classifier performance metric. |
| bvFTD | Behavioural-variant frontotemporal dementia | Patient group. |
| CBI-R | Cambridge Behavioural Inventory-Revised | Behavioural clinical assessment. |
| CON | Healthy control group | Reference group for patient-control contrasts. |
| DCM | Dynamic Causal Modelling | Generative modelling framework used to estimate latent physiological parameters. |
| DMN | Default mode network | Network includes medial prefrontal, posterior cingulate, and angular-gyrus regions. |
| FAB | Frontal Assessment Battery | Frontal-executive clinical assessment. |
| FDR | False-discovery rate | Multiple-comparison control procedure. |
| RAI | Right anterior insula | Fronto-insular/salience-related source region. |
| FRS | Frontotemporal Dementia Rating Scale | Clinical severity scale. |
| LAG | Left angular gyrus | Default-mode/posterior association source region. |
| LOO | Leave-one-out | Cross-validation scheme used for group-to-individual assignment. |
| MCC | Matthew's correlation coefficient | Balanced classifier performance metric robust to class imbalance. |
| MEG | Magnetoencephalography | Resting-state neurophysiological modality analysed in source space. |
| MMSE | Mini-Mental State Examination | Global cognitive assessment. |
| NMDA | N-methyl-D-aspartate receptor | Slower excitatory receptor class represented in DCM parameters. |
| OSL | OHBA Software Library | MEG preprocessing software framework. |
| PAC | Phase-amplitude coupling | Cross-frequency feature linking alpha phase to beta amplitude. |
| PCC | Posterior cingulate cortex | Default-mode source region. |
| PSD | Power spectral density | Frequency-domain representation used to derive empirical spectral features. |
| PSP | Progressive supranuclear palsy | Patient group. |
| PSPRS | Progressive Supranuclear Palsy Rating Scale | Motor and functional clinical severity scale for PSP. |
| RAG | Right angular gyrus | Default-mode/posterior association source region. |
| RBF-SVM | Radial-basis-function support vector machine | Nonlinear classifier used to test curved feature-space separation. |
| SN | Salience network | Network including the left and right anterior insula and anterior cingulate cortex. |
| SPM12 | Statistical Parametric Mapping 12 | Software used for source inversion and DCM procedures. |
| SVM | Support vector machine | Classifier family used for linear and nonlinear assignment. |
